# Whole-neuron synaptic mapping reveals local balance between excitatory and inhibitory synapse organization

**DOI:** 10.1101/395384

**Authors:** Daniel Maxim Iascone, Yujie Li, Uygar Sümbül, Michael Doron, Hanbo Chen, Valentine Andreu, Finola Goudy, Idan Segev, Hanchuan Peng, Franck Polleux

**Author notes:** These authors contributed equally to this work (Daniel Maxim Iascone, Yujie Li).

## Abstract

The balance between excitatory and inhibitory (E and I) synaptic inputs is thought to be critical for information processing in neural circuits. However, little is known about the principles of spatial organization of E and I synapses across the entire dendritic tree of mammalian neurons. We developed a new, open-source, reconstruction platform for mapping the size and spatial distribution of E and I synapses received by individual, genetically-labeled, layer 2/3 cortical pyramidal neurons (PNs) *in vivo*. We mapped over 90,000 E and I synapses across twelve L2/3 PNs and uncovered structured organization of E and I synapses across dendritic domains as well as within individual dendritic segments in these cells. Despite significant, domain-specific, variations in the absolute density of E and I synapses, their ratio is strikingly balanced locally across dendritic segments. Computational modeling indicates that this spatially-precise E/I balance dampens dendritic voltage fluctuations and strongly impacts neuronal firing output.

## INTRODUCTION

The spatial organization of synapses throughout the dendritic tree is a critical determinant of their integration properties and dictates the somatic firing patterns of individual neuronal subtypes (*1-4*). Within dendritic branches, clustered potentiation of excitatory and inhibitory (E and I) synaptic inputs underlie both circuit development and experience-dependent plasticity (*5-9*). Recently there has been substantial progress toward mapping neuronal connectivity at multiple scales (*10-13*). However, significant roadblocks remain in identifying basic principles of synaptic organization for individual neuronal subtypes (*14-17*), leaving important questions unanswered: Are there multiple spatial scales of E and I organization within neurons? Are there hotspots of enhanced synaptic connectivity? Is there a structural correlate of E/I balance within specific dendritic domains or individual dendritic segments? Finally, how does E and I synaptic organization characterizing a given neuronal subtype influence the dendritic and somatic firing properties of these neurons?

Comparing the distributions of excitatory and inhibitory synapses within the same neuron is particularly important to determine the cellular logic of synaptic organization. At the circuit level, a precise balance of excitation and inhibition is critical for calibrating both global and fine-scale levels of activity throughout development and during adult function (*18-20*). In both auditory and somatosensory cortex, the co-tuning of E and I conductances is set by experience-dependent refinement of intracortical inhibition (*21, 22*). An anatomical basis for E/I balance within individual neurons has also been observed in visual cortex and CA1, where excitatory inputs onto pyramidal neurons are continuously offset by somatic inhibition (*23, 24*). A conserved ratio of the numbers of E/I synapses was observed throughout the dendrites of cultured hippocampal neurons, suggesting that the spatial distribution of synapses might also contribute to E/I balance (*2*). However, this finding has not been extended to neurons *in vivo*. Activation of NMDA-type glutamate receptors leads to input-specific long term potentiation of dendritic inhibition mediated by somatostatin-expressing interneurons, linking excitation and inhibition within individual dendritic segments (*25*).

Here we have developed an adaptable, open-source platform for imaging and mapping E and I synapses across the entire dendritic arbor of individual neurons. We created whole-cell reconstructions of individual, optically-isolated pyramidal neurons (PNs) containing information about the size, shape, and continuous position of all E and I synapses across their entire dendritic arbors: the first dataset of its kind for any neuronal subtype. We focused our study on layer 2/3 (L2/3) PNs of the adult mouse primary somatosensory cortex, where substantial prior knowledge of the synaptic microstructure and connectivity allowed validation of our platform and some of our findings (*26-31*) as well as identification of new principles of E and I synaptic organization.

## RESULTS

### Synapse Detector: a Platform to Create Whole-Neuron Structural Inputs Maps

To obtain optically isolated, single L2/3 PNs for these synaptic reconstructions, we co-electroporated Cre-dependent Flex-tdTomato with low levels of Cre recombinase for extremely sparse *in utero* electroporation ((*32, 33*); **Fig. 1A** and **Movie S1**). We also labeled inhibitory synapses received by individual PNs by co-electroporating the inhibitory postsynaptic scaffolding protein Gephyrin tagged with EGFP, a strategy previously shown to reliably label all GABAergic and glycinergic inputs without affecting their development (*9, 34*). We achieved single-synapse resolution using confocal microscopy by imaging neurons across 2-3 serial 150 μm vibratome sections with a 100x 1.49 NA objective lens (**Fig. 1B** and **1D** and **Movie S2**).

**Figure 1.**
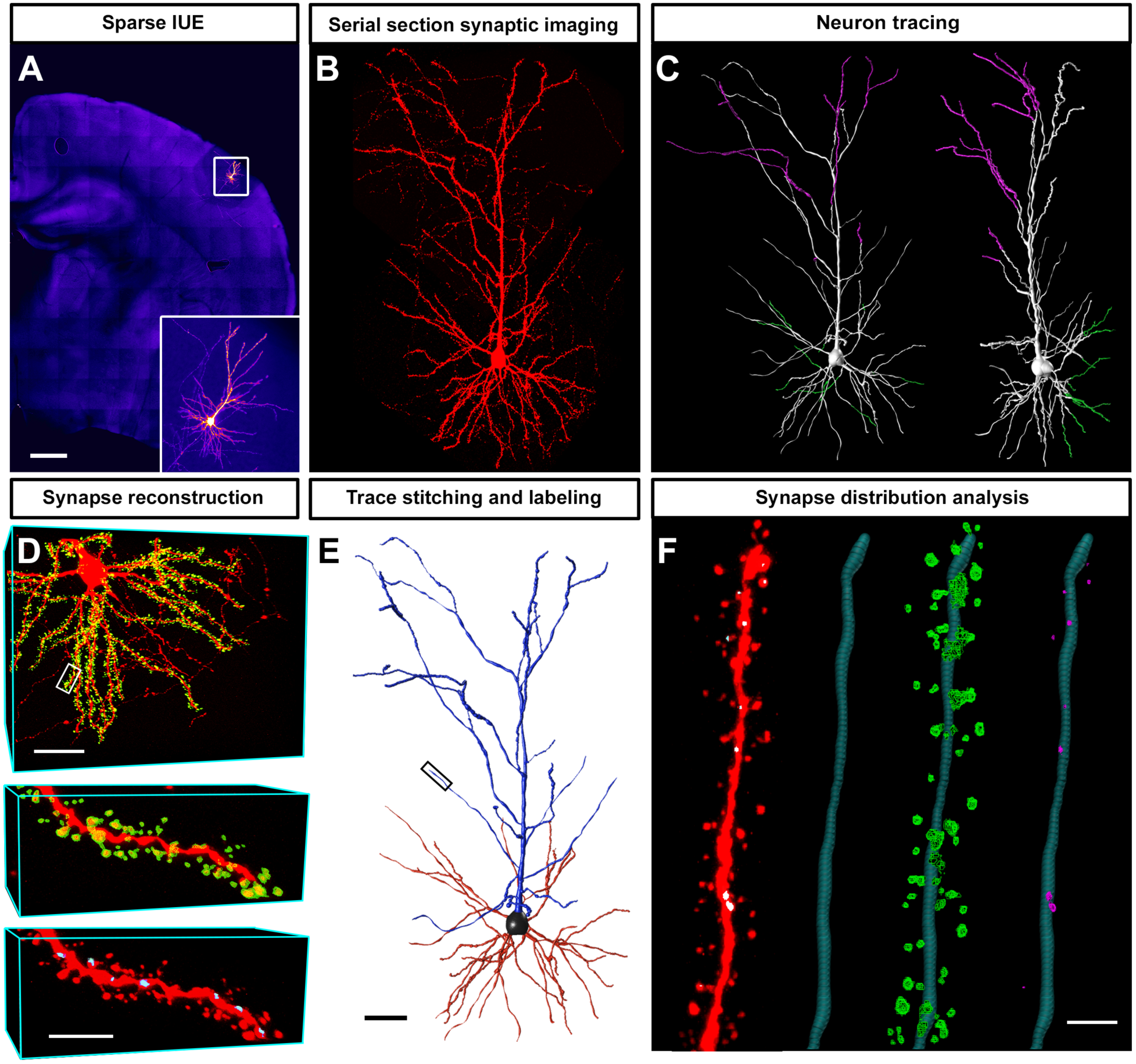
Spatial and morphological annotation of synapses across whole neurons with Synapse Detector. **(A)** Sparsely labeled layer 2/3 pyramidal neuron (PN). Scale bar: 500 microns. Inset, higher magnification of neuron shown in **A**. **(B)** Single-synapse resolution image volume compiled from 3 adjacent sections containing a complete layer 2/3 PN expressing tdTomato and Gephyrin-eGFP. **(C)** Neuron trace of **B** (left) and rotated to display the tissue section plane (right). Each color (Magenta, silver and green) represents parts of the dendritic arbor traced and stitched from individual adjacent sections. **(D)** Dendritic spines annotated throughout the basal dendritic arbor of **B** (top panel). Scale bar: 50 microns. Annotated spines (green in middle panel), and inhibitory Gephyrin-eGFP labeled synapses contained (blue in bottom panel) from top panel inset. Scale bar: 5 microns. **(E)** Neuron trace of **B** annotated with Subtree Labeling program. Scale bar: 50 microns. **(F)** From left to right: tdTomato volume from inset in **E**, corresponding trace, overlaid spine annotations, and inhibitory synaptic annotations associated with nodes within the trace. Scale bar: 2 microns.

This new Synapse Detector toolkit within Vaa3D generates synaptic maps by taking image data and a trace of the dendritic tree as input to automatically isolate E and I synapses within a user-defined radius of each dendrite. Within this toolkit, excitatory synapses (dendritic spines) are classified with a Spine Detector module that identifies regions of fluorescence surrounding the dendritic trace (**Fig. 1D** top and middle panels and **Movies S3-S5**). Inhibitory synapses are identified using an IS Detector program that identifies EGFP-Gephyrin puncta that co-localize with the cytosolic Flex-tdTomato (**Fig. 1D** bottom panel and **Movies S6** and **S7**). Together these software platforms measure E and I synapse position in 3D along the dendritic tree, as well as morphological features of E and I synapses such as their volume, spine neck length, position of I synapses along dendritic shaft or on spine heads (so called dually innervated spines, (*9, 27, 35*)). During reconstruction, Synapse Detector’s editing features allow the user to edit the volume of each identified synapse and eliminate false positives (**Fig. S1**). Synapse Detector has a minimal false negative rate compared to manual reconstructions and generates consistent annotation results among multiple users (**Fig. S3**). Following reconstruction and manual annotation, synaptic features are associated with nodes providing their geometric position in 3D along the dendritic tree. In the final step, neuron trace fragments containing information about individual synapse position and size from serial tissue sections are stitched together into a final input map with the Vaa3D Neuron Stitcher program that can be used to analyze the morphology of all synapses as a function of their continuous distance from the soma along the dendritic arborization (*36*).

We used this new Vaa3D reconstruction pipeline to map all E and I synapses across 10 PNs from L2/3 primary somatosensory cortex (as well as excitatory synapses from 2 additional PNs; **Fig. 2**). These neurons contained on average 6773±212 dendritic spines (range: 5558-8115) and 939±101 inhibitory synapses (range: 595-1556; **Fig. 2C**). On average, the total length of these dendritic trees was 4579±103 µm and these neurons displayed overall E and I synaptic densities consistent with previous reports (1.48±0.04 spines/µm and 0.20±0.02 inhibitory synapses/µm respectively) ((*9, 35, 37*); **Fig. 2C**). We also found that 26%±2% of inhibitory synapses targeted dendritic spines in L2/3 PNs (**Fig. 2C**). This fraction of spines dually innervated by an E and I synapse is comparable with previously observed values in these neurons (*9, 27, 35*).

**Figure 2.**
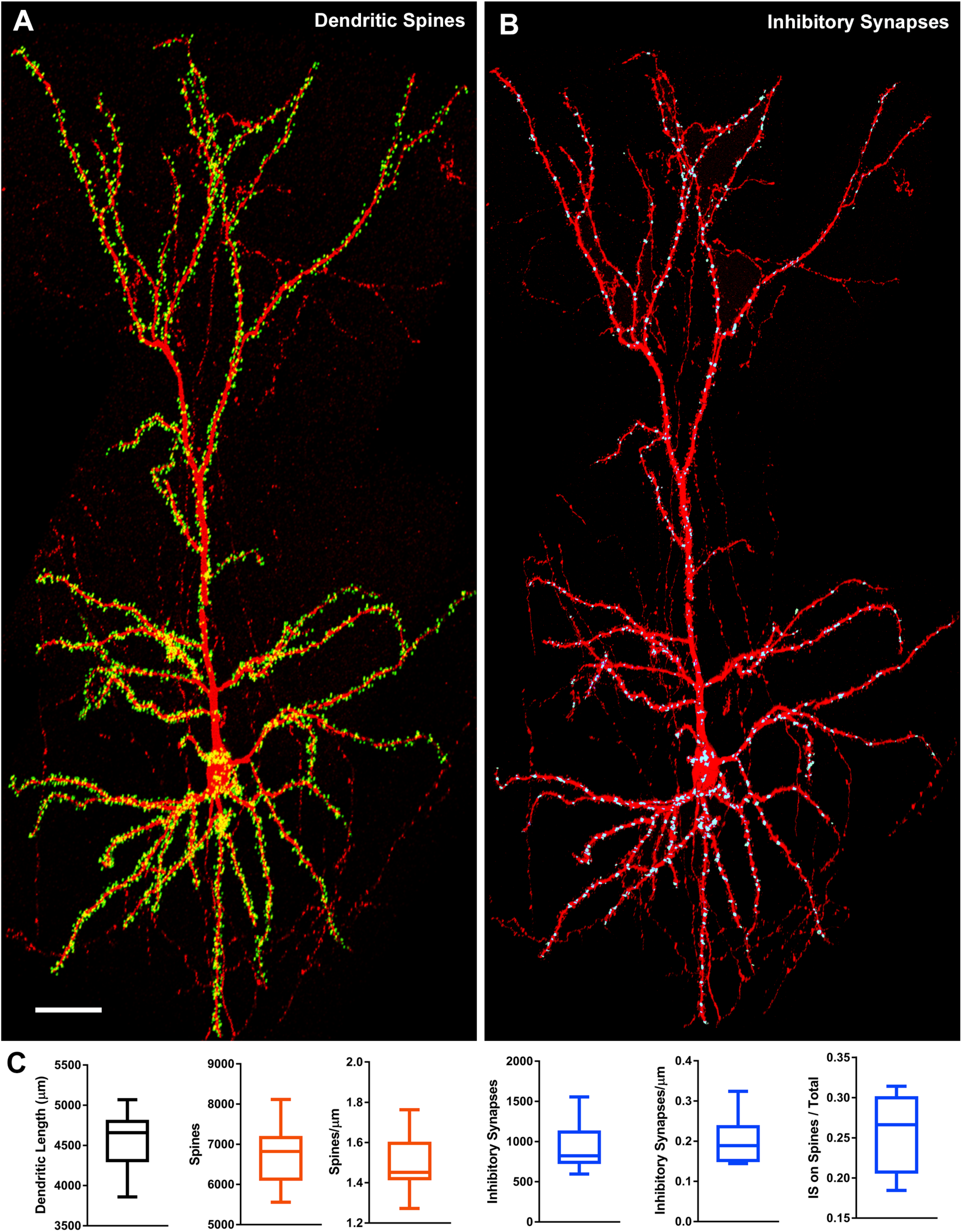
Synaptic distribution profile for layer 2/3 somatosensory PNs. **(A-B)** Example of complete reconstruction of 8115 dendritic spines (green in A) and 1045 I synapses (blue in B) throughout the dendritic tree of a single layer 2/3 PN (Neuron 1; red cell-filler, tdTomato). **(C)** From left to right: box plots showing the distribution of dendritic length, dendritic spine number, dendritic spine density, inhibitory synaptic number, inhibitory synaptic density, and dually innervated spine proportions for all layer 2/3 PNs mapped in this study. Scale bar: A-B, 50 microns.

### Features of E and I synaptic organization across the entire dendritic tree of layer 2/3 PNs

To analyze synaptic distribution across the entire dendrites of the reconstructed L2/3 PNs, we subdivided dendritic arbors into three distinct domains: apical tuft, apical oblique, and basal dendrites (*38*). Within these domains, we distinguished among segment types by their relative branch order: primary, intermediate, and terminal ((*38*); **Fig. 3A**). This categorization is functionally relevant as different branch orders have distinct passive conductance properties resulting from their relative size and distance to the soma (*38, 39*). Primary dendrites have relatively low input impedance due to their large size, while terminal dendrites have higher input impedance due to their smaller diameter and sealed end. In addition to the domain classification used here, we developed a Subtree Labeling program as part of the Spine Detector toolkit that enables user-directed annotation of regions of interest throughout the neuron trace to assess experiment-specific questions about domain-level synaptic organization (**Fig. S2E** and see **Material and Methods**).

**Figure 3.**
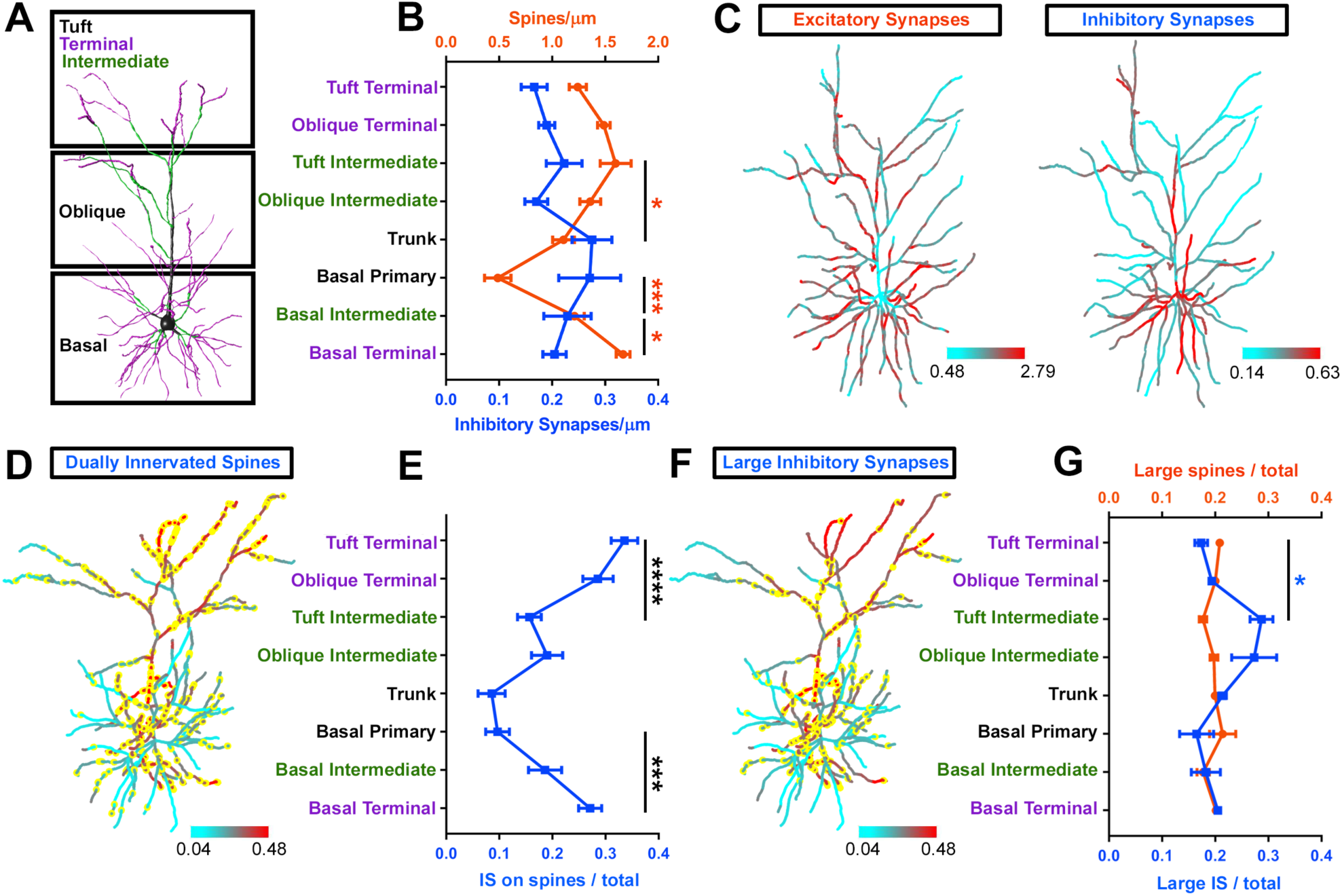
Domain organization of synaptic distribution and morphology. **(A)** A schematic diagram of a L2/3 PN depicting the domains (black boxes) and branch types (black primary; green, intermediate; and purple, terminal). **(B)** The density of E (orange line) and I (blue line) synapses across dendritic branch types of L2/3 PNs. **(C)** Heat maps of E (left) and I (right) synaptic distribution indicating regions of low density in cyan and high density in red. **(D)** A heat map of inhibitory synaptic distribution in which yellow puncta represent inhibitory synapses targeted to dendritic spines. Note the increased density of these dually innervated spines toward the distal apical tufts. **(E)** The proportion of inhibitory synapses made onto dendritic spines across dendritic branch types. **(F)** A heat map of inhibitory synaptic distribution in which yellow puncta represent the 20^th^ percentile of inhibitory synapses by volume for each domain type. Note the increased density of these large inhibitory synapses in apical intermediate segments. **(G)** The proportion of large excitatory (orange) and inhibitory (blue) synapses across dendritic branch types. For all plots, *p < 0.05, **p < 0.005, and ***p < 0.001. See **Material and Methods** for details. All data are presented as mean ± SEM.

This division of the dendritic tree into specific domains and branch types allowed us to characterize the profile of synaptic distribution across L2/3 PNs. Similar to previous observations in CA1 PNs, E and I synaptic distribution appear to be inversely correlated at the domain level with relatively low spine density proximal to the soma, suggesting that this may be a general feature of synaptic organization across PN subtypes ((*40, 41*); **Fig. 3B**). In contrast to previous studies however, our complete reconstructions enable whole-cell mapping of relative E and I synaptic distribution (**Fig. 2C**, **S4** and **S5**). Maps of E and I synaptic distribution in the same neuron demonstrate an almost complete absence of spines along primary dendrites accompanied by the highest density of inhibitory synapses (**Fig. 3C**).

L2/3 PNs receive direct thalamic input from both the ventral posteromedial nucleus (VPM) and the posterior medial nucleus (POm) terminating mainly onto their apical tufts (L1) and basal dendrites (L3) respectively (*28, 42*). The vast majority of neocortical spines that are dually innervated by an inhibitory synapse receive excitatory inputs from corticothalamic axons, suggesting that distal tuft and basal dendrites should have a relatively high proportion of dually innervated spines (*27*). Our unbiased mapping of the location of IS located on spine heads demonstrates that apical tuft and basal terminal dendrites of L2/3 PNs display a significantly higher proportion of dually innervated spines than primary and intermediate dendrites (**Fig. 3D-E**), validating the spatial resolution of our labeling, imaging, and reconstruction approaches.

Because spine head volume is linearly proportional to excitatory synaptic strength (e.g. size of the post-synaptic density and density of glutamatergic AMPA receptors), it is also possible to use Synapse Detector to map the distribution of relative synaptic strengths (*3, 43, 44*). We classified “large” synapses as greater than the highest 20^th^ percentile of synaptic volume for each neuron, closely corresponding to the persistent 160% increase in volume reported for synapses following structural forms of long-term potentiation (*5, 45, 46*). Interestingly, while there is no specific trend for the distribution of large spines across dendritic domains, large inhibitory synapses appear to be clustered around the apical intermediate dendritic segments (**Figures 3F** and **3G**), a feature never detected before.

### E and I synaptic distribution is structured and locally balanced in dendritic segments

Active dendritic conductances evoked by clustered synaptic inputs can produce nonlinear depolarization and change the probability of somatic firing (*47*). Local increases in excitatory synaptic density in a subset of segments within the same dendritic domain could reflect clustered spine stabilization following branch-specific synaptic potentiation (*5, 47*). Therefore, we tested if L2/3 PNs exhibit local changes in the relative distribution of E and I synapses across segments within each dendritic domain (**Fig. 4**). To assess the extent of this potential weighted synaptic distribution, we compared the experimentally observed variation in synaptic density between segments within each dendritic domain to randomly shuffled densities for each neuron reconstructed. This was done by randomly redistributing synaptic density values across segments of the same domain (see Supplementary Material). Neurons in which synaptic distribution is significantly weighted toward a subset of dendritic segments would therefore display greater domain-specific variation in synaptic density than correspondingly randomized versions. Excitatory synaptic (spine) distribution is significantly weighted toward a subpopulation of dendritic segments across almost the entire dendritic tree (**Fig. 4A** and **4D**). Interestingly, inhibitory synaptic distribution is significantly non-random and clustered only in apical and basal terminal domains, raising the intriguing possibility that E and I synapses are weighted toward the same dendritic segments (**Fig. 4B** and **4D**).

**Figure 4.**
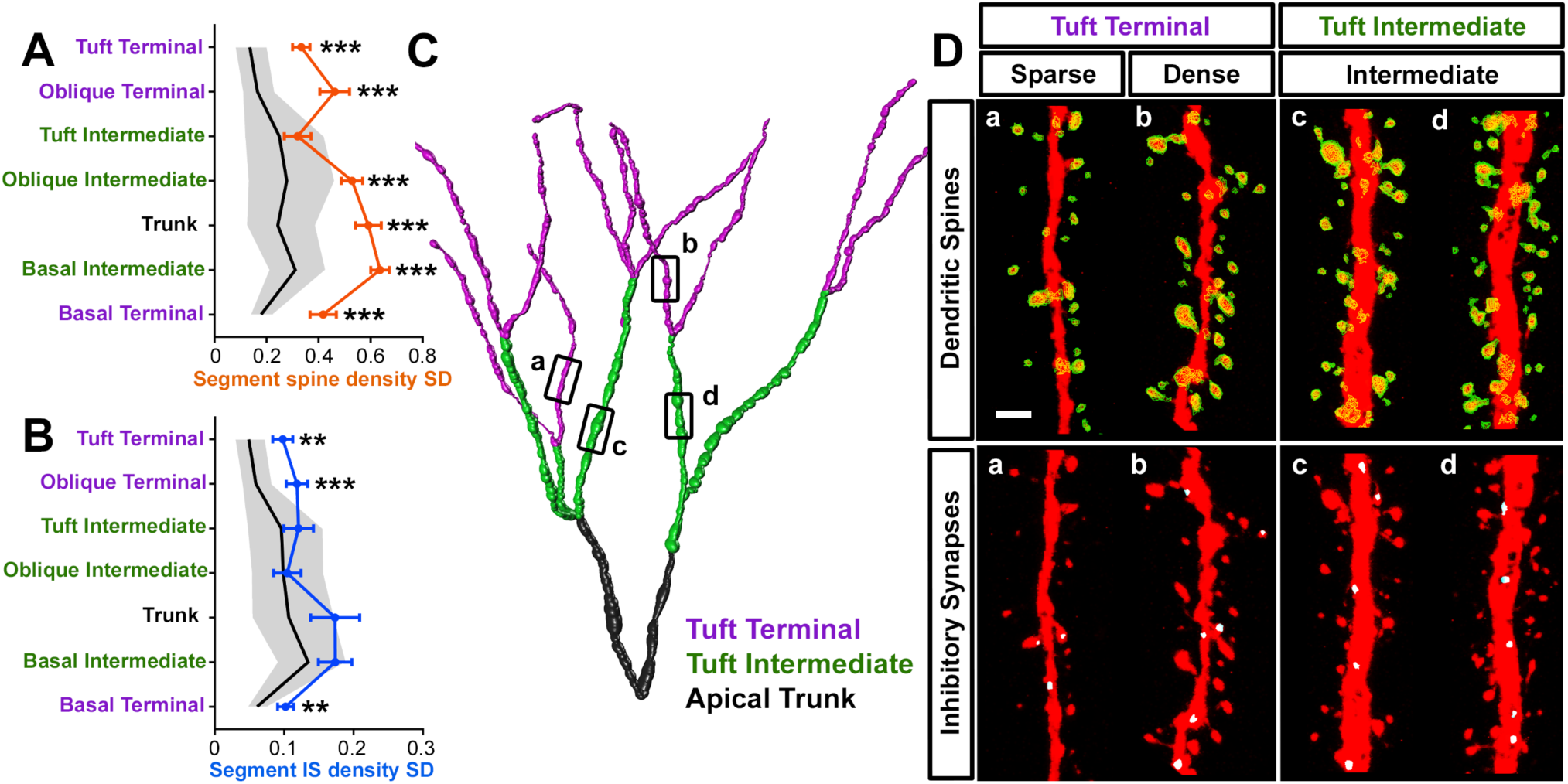
Structured synaptic distribution within branch types. **(A)** Variation in excitatory synaptic density (orange) across dendritic branch types of L2/3 PNs compared to 10,000 randomizations of each synaptic distribution within the same domains (5^th^ to 95^th^ percentiles gray). **(B)** Variation in inhibitory synaptic density (blue) across dendritic branch types of L2/3 PNs compared to randomized synaptic distributions (gray). **(C)** Example trace of apical tuft dendrites depicting the breakdown of the arbor into branch types (trunk: black, intermediate: green, and terminal: purple). **(D)** Excitatory (top) and inhibitory (bottom) synapses from segments isolated from the dendritic arbor in **C**. Terminal tuft segments (left) display significant variation in E and I synaptic density while intermediate tuft segments (right) do not. Scale bar: 1 micron.

Co-regulation of E and I synaptic inputs, generally referred to as E/I balance, is a critical mechanism for calibrating both global and fine-scale levels of neuronal activity (*23, 48, 49*). While several studies have demonstrated mechanistic links between E and I synaptic potentiation, whether it results in local, fine-scale balance between E and I synaptic distribution within dendritic segments remains an open question (*25, 46, 50*). Remarkably, we find that E and I synaptic density strongly co-varied in terminal segments throughout the dendritic tree of layer 2/3 PNs (**Figure 5**). This structural E/I balance appears to increase as a function of distance from the soma, with segments distal to the soma showing remarkable correlation between E and I synaptic density (**Figure 5**). Taken together, these results demonstrate that in L2/3 PNs: (1) both E and I synaptic density varies more than by chance between dendritic segments among a given dendritic domain and (2) that despite this variability in E and I synapse density between segments of a given dendritic domain, the ratio between E and I synaptic density is tightly controlled locally within these segments.

**Figure 5.**
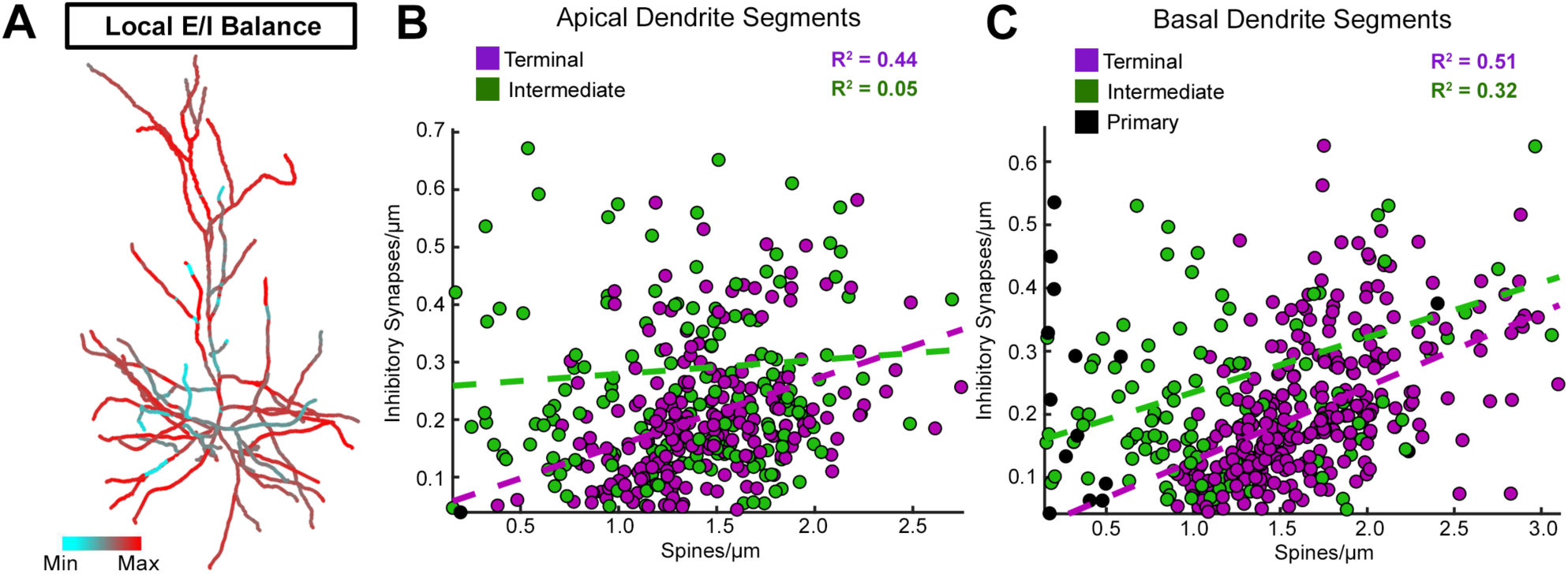
Local E/I balance within branch types. **(A)** Heat map of local E/I balance. Densities of E and I synapses were normalized to 1, and the absolute value of their difference is mapped within an adaptive range of each point within the dendritic tree (see **Material and Methods**). Values close to 0 therefore represent points on the dendritic tree at which the relative densities of E and I synapses were close to equivalent (Max), and values close to 1 represent points at which the relative densities of E and I synapses were much different from one another (Min). **(B)** Relation between E and I synaptic density for intermediate (green; R^2^ = 0.05; p > 0.05) and terminal (purple; R^2^ = 0.44; p < 1.82e^-27^) apical dendritic segments. **(C)** Relation between E and I synaptic density for primary (black), intermediate (green; R^2^ = 0.32; p < 0.0003), and terminal (purple; R^2^ = 0.51; p < 2.95e^-21^) basal dendritic segments. For all plots, *p < 0.05, **p < 0.005, and ***p < 0.001. See **Material and Methods** for details. All data are presented as mean ± SEM.

### Functional implications of global and domain-specific E/I balance

To better understand the functional implications of the local E/I balance we found experimentally (**Figure 5**), we performed computational modeling of the 10 individual L2/3 PNs reconstructed (shown in **Fig.S4 and S5)**, including their 3D reconstructed morphology and the dendritic location of their E and I synapses. Passive and active membrane properties of these cells were based on previously published biological values (see **Material and Methods**). All excitatory synapses were activated randomly at an average rate of 1.75 HZ while inhibitory synapses were activated at 10 Hz so that the firing rate of the modeled cells matched ranges found *in vivo* (*31*). These models also replicated several active and passive dendritic properties observed in L2/3 PNs, including back-propagating action potentials, the somatic input resistance, and membrane time constants (**Figure 6** and see **Material and Methods**).

**Figure 6.**
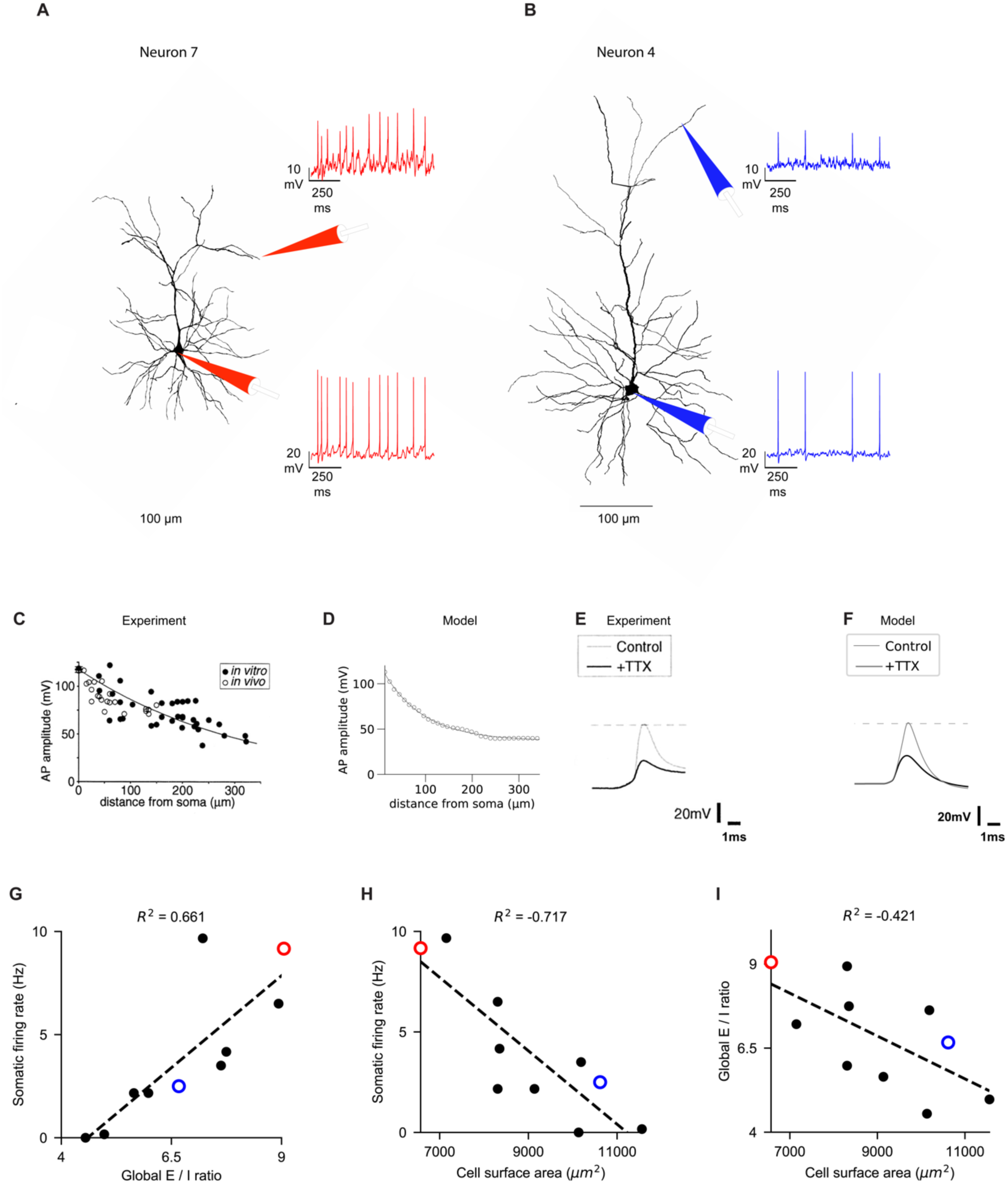
Models predict large variability in the firing rate of L2/3 PNs. L2/3 neurons with larger dendritic membrane area tend to have smaller E/I ratios, leading to a lower firing rate. **(A)** The reconstructed morphology of Neuron #7. Simulated synapses were placed on the experimentally-measured locations of the spines (excitatory synapses, 5,604 in total) and of the inhibitory synapses (619 in total) and activated randomly as in **Fig. 4**. Simulated membrane voltage was recorded from a distal apical dendrite (upper trace) and from the soma (lower trace). **(B)** As in **A**, with the morphology and traces belonging to Neuron #4 (5,661 excitatory synapses and 785 inhibitory synapses). **(C)** Backpropagating action potentials, BPAP) in L2/3 PNs attenuate along the apical trunk (taken from the experiments of (*60*)). Y-axis shows the amplitude of the BPAP, measured at different distances from the soma along the apical trunk (x axis), both *in vivo* (empty dots) and *in vitro* (solid dots). **(D)** Same as in **C**, in the simulated L2/3 model. **(E)** Action potential amplitude recorded 80 µm from the soma in the apical trunk, with and without TTX (from (*60*)). **(F)** Same as in **E**, in L2/3 model. **(G)** The correlation between somatic firing rate and the global E/I ratio for all 10 modeled neurons; filled circles are for different cells. Pearson coefficient is shown above the graph. Red and blue dots correspond to neurons #7 and #4 shown in **A** and **B**, respectively. **(H)** Same as in **G**, with the x-axis showing the cell’s surface area. **(I)** Global E/I ratio as a function of the total dendritic surface area.

To test the significance of the synaptic distribution we observed in these 10 L2/3 PNs, we manipulated the variance (here measured as standard deviation, SD) of the domain-specific E/I ratio while keeping the total number of synapses within each domain constant (thus keeping the global E/I ratio fixed for the modeled cell). This created a range of E/I ratio SD values, ranging from very tight E/I ratio variance, where each segment within a given domain has a similar ratio of E and I synapses (**Fig. 7A**, *the balanced case*, left), to the extreme case, in which each branch in a given domain had either only excitatory or only inhibitory synapses (**Fig. 7B**, *the unbalanced case*, left). Manipulation of the variance of segment-specific E/I ratio had a strong effects on predicted dendritic voltage dynamics (**Fig. 7A-B**, right): variation in dendritic voltage (including active dendritic spiking and back propagating action potentials (Material and Methods in **Fig. 7A-7B**) in balanced segments is dampened and overall more hyperpolarized (**Figure 7A**, right) compared to the unbalanced E and I cases (**Fig. 7B**, right). In the case of minimal variance of local E/I ratio per segment, the voltage distribution was narrower and very similar to that predicted from the biologically-observed E/I ratio (compare blue to green lines in **Fig.7C**). We found that terminal domains which, experimentally, had a near-balanced E/I ratio (**Fig. 5**) were highly sensitive to increasing the E/I ratio variance: gradually increasing the variance of E/I ratio among segments resulted in a gradual increase of the mean dendritic voltage time-integral (**Fig. 7D**). This was not the case for intermediate domains with biologically unbalanced E/I ratio, where the change in variance of E/I ratio between segments had minimal effect on the dendritic voltage time-integral (**Fig. 7G** and see **Material and Methods**). This is likely due to the small contribution of intermediate synapses to the voltage perturbations in those domains, possibly as a result of their small relative number: indeed, if we perform our simulations after removing the synapses in intermediate domains, of the voltage integral was reduced by less than 1%, compared to 10-30% in the terminal domains (**Fig. S7A**).

**Figure 7.**
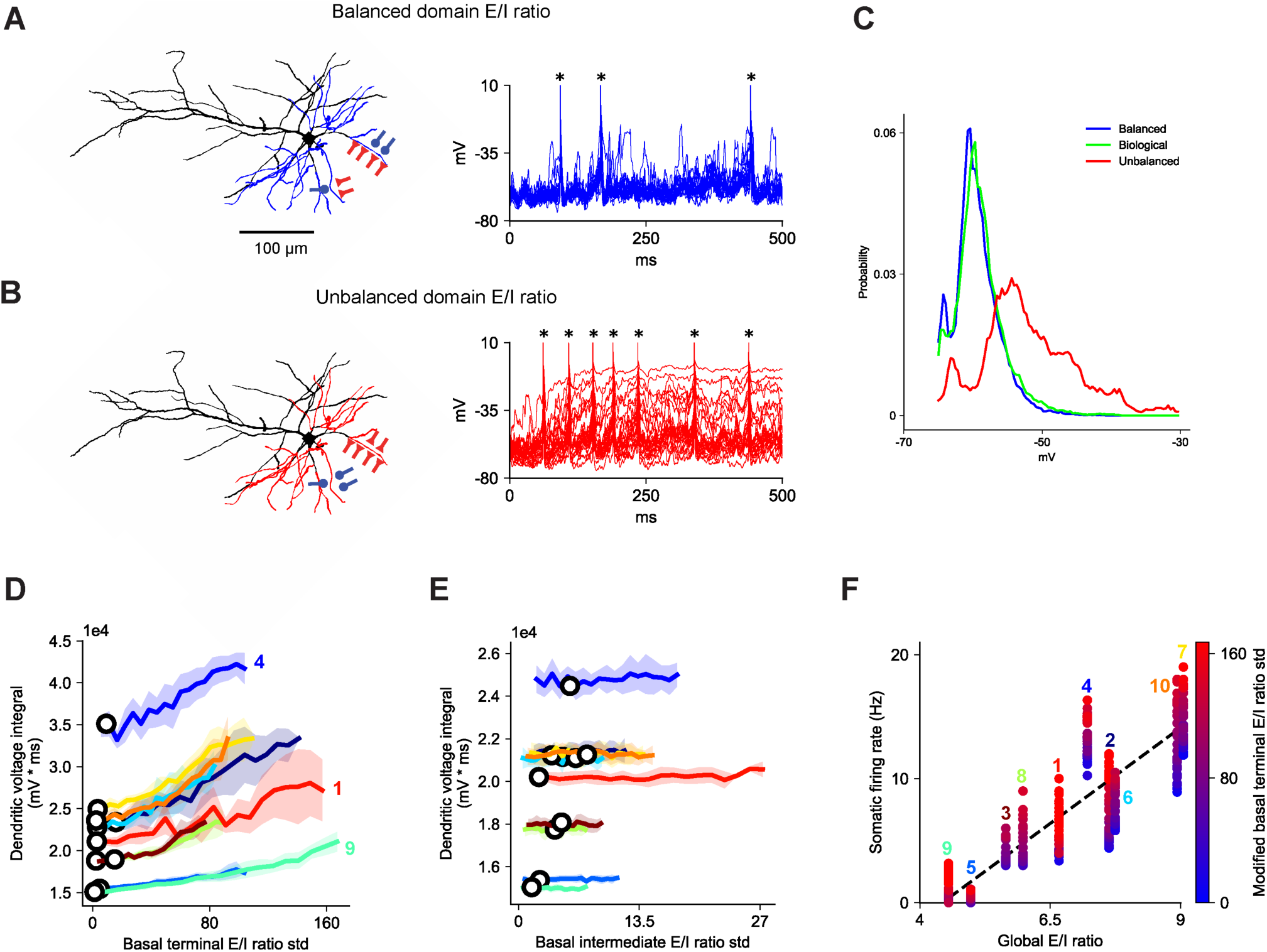
Domain-specific E/I balance within L2/3 PNs dampens local dendritic voltage fluctuations, strongly affecting the global output firing rate. **(A)** (Left) The modeled L2/3 cell (#2) for the case depicted schematically, in which the E/I ratio is constant for all branches belonging to the distal apical domain (zero E/I variance, the *balance case*). (Right) Voltage traces in the various distal apical branches for the balanced case in response to activation of all excitatory synapses (6,274 in total) and inhibitory synapses (1,111 in total) as measured experimentally for this cell. Back propagating action potentials are marked by an asterisk. **(B)** Same as in **A** for maximal E/I SD (the *unbalanced case*). In both **A** and **B,** the total number of E and I synapses in the distal apical domain is fixed as found experimentally (same global balance for the two cases). Note that back-propagating action potentials (large depolarizing transient) exhibit higher frequencies in the unbalanced case. **(C)** The probability of voltage integral for all branches in the basal terminal domain at the modeled cell. The distribution of the dendritic voltage time-integral expected for the experimentally-measured case (green line) closely fits that expected in the balanced case (blue line); both are narrower and less depolarized as compared to that obtained in the unbalanced case (red line). **(D)** The average voltage time-integral for all segments in the basal terminal domain as a function of E/I SD in this domain (see **Material and Methods**). The open circles represent the biologically measured E/I SD values for each neuron. Numbers correspond to cell numbers in **Figure S4**. **(E)** As in **D** for the basal intermediate segments. **(F)** Correlation between the somatic firing rate and the global E/I ratio for the 10 modeled cells. In each case (numbered vertical lines), the domain-specific E/I SD varied from the fully balanced case (blue) to the maximally unbalanced case (red). Numbers at each vertical line correspond to the modeled cell identity. In all cases, the numbers for activated E and I synapses were taken from the experimental counts; excitatory synapses were all activated randomly with an average rate of 1.75 Hz and inhibitory synapses were activated at 10 Hz. See details in **Material and Methods**.

The somatic firing rate in the 10 modeled cells was also strongly affected by the domain-specific E/I SD value (keeping the global E/I balance fixed per cell). Indeed, when testing the combined effect of differences in the global E/I ratio for different modeled cells together with the effect of the domain-specific E/I SD, we found a high correlation (R^2^ = 0.7, dashed line) between somatic firing rate and global E/I ratio (**Figure 7F**). Not surprisingly, cells with larger relative number of excitatory synapses fire at higher rates. Strikingly, in all modeled cells, the output firing rate increases as much as twofold per cell when the domain specific E/I ratio SD was increased, suggesting that the local E/I ratio (in addition to the global E/I ratio) must be considered for understanding how synaptic activity shapes the neuron’s output. Our experimentally-based modeling demonstrates that in L2/3 PNs, E/I ratio is optimally-balanced in key dendritic domains; this domain-specific, local E/I balance at the level of dendritic segments constrains dendritic voltage fluctuations, and controls to a significant extent the firing rate of these neurons (**Figures 7F** and **S6**).

## DISCUSSION

Mapping the spatial organization of synapses across the entire dendritic arbor of individual neurons is crucial for bridging the gap between our understanding of the molecular determinants of synaptic development and the principles of neural circuit connectivity. Here, we developed an adaptable, open-source toolkit for mapping the morphology and spatial distribution of all E and I synapses across complete neurons. This method has several key benefits for mapping subcellular synaptic morphology and distribution. As part of the Vaa3D image annotation platform, Synapse Detector is fully integrated into Vaa3D automatic pipeline for image segmentation, 3D image stitching, and surface reconstruction (*36, 51, 52*). Synapse Detector is compatible with any fluorescent imaging method including high resolution confocal microscopy. Most importantly, Synapse Detector provides a generalizable toolkit for quantifying and mapping features of subcellular fluorescent marker distribution.

The synaptic mapping pipeline developed here enabled the reconstruction of 12 L2/3 PNs, including the location and morphology of over 90,000 E and I synapses. Previous anatomical studies of the synaptic morphology and connectivity of this cell type allowed validation of our platform and observed results (*26-31*). Our 3D reconstruction method for dendritic spines closely matched estimates from manual reconstructions of L2/3 PN spine density and morphology generated by tracing synapses from serial focal planes (*26*). We also observed inhibitory synaptic distributions consistent with previous observations, as well as a common proportion of inhibitory synapses targeted to spines (*9, 34, 35*). Strikingly, the distribution of specific synaptic features characterizing mouse L2/3 PNs recapitulates known motifs of circuit connectivity: in L2/3 PNs, dually innervated spines almost exclusively correspond to dendritic spines receiving thalamic inputs, and these synapses were significantly enriched in the L1 apical tufts and deep L3 basal dendrites, the two layers targeted by thalamic afferents from POm and VPL that innervate S1 (*27, 28*).

Analyzing the distribution of E and I synapses across complete dendritic arbors has revealed several scales of structured organization within L2/3 PNs. E and I synaptic distribution also varies significantly across dendritic domains, with fewer spines located in proximal than along distal dendritic segments, similar to what has been observed in CA1 PNs, potentially suggesting a shared principle for synaptic organization between these PN subtypes (*40, 41*).

Crucially, within-neuron comparisons of observed and randomized synaptic locations enabled the identification of structured distribution of E and I synapses to a restricted subset of dendritic segments within each domain. The formation of hotspots of synaptic density is consistent with known cellular mechanisms promoting spatially clustered synaptic stabilization and potentiation at the scale of single dendritic segments (*47, 53*). Active properties of dendrites critical for initiating clustered potentiation are engaged in somatosensory and visual cortical PNs during sensory processing, raising the tantalizing possibility that these hotspots of synaptic density might represent a structural signature of salient feature storage within neuronal dendrites (*8, 54-56*).

A novel feature of structured synaptic distribution that emerged from our study is the strong, branch-specific, and local balance between E and I synaptic density across terminal dendritic segments. This suggests a far stronger association between E and I synaptic distribution than previous observations *in vitro* that the total number of E and I synapses are correlated across dendrites (*2*). Indeed, while a conserved ratio of the number of E and I inputs across dendrites can be largely explained by longer dendrites receiving more inputs, our data strongly suggest that molecular mechanisms co-regulating the balance between E and I synaptic density must be acting at the scale of short dendritic segments. This spatial pattern closely matches the dendritic targeting of somatostatin-expressing interneurons, whose synapses onto L2/3 PNs were recently demonstrated to undergo NMDAR-dependent long term potentiation (*25, 57, 58*). While the study of E/I balance at the level of single neurons has largely been restricted to feedforward inhibition mediated by perisomatic-targeting basket interneurons, our whole-neuron synaptic input maps suggest that a precise balance between excitation and inhibition is critical for dendritic integration as well (*23, 59*).

This novel principle of local E/I balance within dendritic segments has significant implications for dendritic integration. Indeed, our simulations show that disrupting the biologically-observed dendritic E/I balance in terminal dendrites dramatically enhances local dendritic voltage fluctuations and the initiation of local dendritic non-linearities, resulting in increased firing at the soma. Our first-ever complete mapping of E and I synapses over the whole dendritic tree of a subtype of PNs, combined with detailed simulations, suggest that the fine-tuned spatial balance of E and I synapses we observed strongly impacts local dendritic computation as well as the global input/output dynamics of cortical neurons within a network. Finally, we provide here the open-source synaptic reconstruction tools we have developed as well as our complete data set of 12 pyramidal neuron input maps containing information about the size, shape, and placement of over 90,000 E and I synapses publicly available upon acceptance.

## ACKNOWLEDGEMENTS

We thank Attila Losonczy, Wes Grueber, Inbal Israeli, Larry Abbott and members of the Polleux lab for fruitful discussions. This work was supported by grants from the NIH (RO1 NS067557 to F.P. and F31 NS101820 to D.M.I.), the NSF (1564736 to Y.L.), ARO MURI (W911NF-12-1-0594 to U.S.), and DoI/IBC IARPA (D16PC00008 to U.S.). I.S. was supported by grant agreement no. 604102 ‘Human Brain Project’ and by a grant from the Gatsby Charitable Foundation. Contributions: D.M.I. and F.P. designed the study. D.M.I. developed the labeling/imaging protocol and performed cloning, animal surgery, and imaging. Y.L. and H.P. developed the Synapse Detector and Subtree Labeling programs with assistance from D.M.I. for user interface design. U.S. developed the anatomical synaptic distribution analysis with assistance from D.M.I. and generated neuron heat maps. M.D. and I.S. performed computational modeling experiments in coordination with D.M.I. and F.P. H.C. developed the trace stitching program. D.M.I., V.A., and F.G. generated neuron reconstructions using Synapse Detector. D.M.I., Y.L., U.S., M.D., I.S., H.P., and F.P. wrote the paper.

## MATERIALS AND METHODS

### Principles of excitatory and inhibitory synaptic organization constrain dendritic spiking in pyramidal neurons

Iascone and Li et al.

### This PDF file includes

Key Resources Table Figures S1 to S7 Methods

Synapse Detector User’s Guide References

Other Supplementary Material includes Movies S1 to S7 and 12 L2/3 PN synaptic maps

**KEY RESOURCES TABLE**

| REAGENT or RESOURCE | SOURCE | IDENTIFIER |
| --- | --- | --- |
| Critical Commercial Assays |  |  |
| In-Fusion HD cloning kit | Clontech | 639649 |
| Deposited Data |  |  |
| P42 CD-1 IGS mouse L2/3 S1 neuron 1 (E and I) | This Paper | N/A |
| P42 CD-1 IGS mouse L2/3 S1 neuron 2 (E and I) | This Paper | N/A |
| P42 CD-1 IGS mouse L2/3 S1 neuron 3 (E and I) | This Paper | N/A |
| P42 CD-1 IGS mouse L2/3 S1 neuron 4 (E and I) | This Paper | N/A |
| P42 CD-1 IGS mouse L2/3 S1 neuron 5 (E and I) | This Paper | N/A |
| P42 CD-1 IGS mouse L2/3 S1 neuron 6 (E and I) | This Paper | N/A |
| P42 CD-1 IGS mouse L2/3 S1 neuron 7 (E and I) | This Paper | N/A |
| P42 CD-1 IGS mouse L2/3 S1 neuron 8 (E and I) | This Paper | N/A |
| P42 CD-1 IGS mouse L2/3 S1 neuron 9 (E and I) | This Paper | N/A |
| P42 CD-1 IGS mouse L2/3 S1 neuron 10 (E and I) | This Paper | N/A |
| P42 CD-1 IGS mouse L2/3 S1 neuron 11 (E only) | This Paper | N/A |
| P42 CD-1 IGS mouse L2/3 S1 neuron 12 (E only) | This Paper | N/A |
| Experimental Models: Organisms/Strains |  |  |
| Mouse: CD-1 IGS | Charles River | Cat# 022 |
| Recombinant DNA |  |  |
| Plasmid: pCAG NLS-Cre | This Paper | N/A |
| Plasmid: pCAG EGFP-GPHN | This Paper | N/A |
| Plasmid: pEf1α FLEX-tdTomato | This Paper | N/A |
| Software and Algorithms |  |  |
| Vaa3D | Peng et al., 2010 | vaa3d.org/ |
| Spine Detector | This Paper | vaa3d.org/ |
| IS Detector | This Paper | vaa3d.org/ |
| Subtree Labeling | This Paper | vaa3d.org/ |
| Neuron Stitcher | Chen et al., 2017 | vaa3d.org/ |
| NEURON | Carnevale, N.T. and Hines, M.L. The NEURON Book. Cambridge, UK: Cambridge University Press, 2006. | https://www.neuron.yale.edu/ |
| MATLAB | The MathWorks Inc. | http://ch.mathworks.com/products/matlab/ |
| GraphPad PRISM | GraphPad Software Inc. | http://www.graphpad.com/scientific-software/prism/ |
| Other |  |  |
| A1R confocal microscope | Nikon | https://www.nikoninstruments.com/Products/Confocal-Microscopes/A1R-HD |

## SUPPLEMENTAL FIGURES

**Figure S1.**
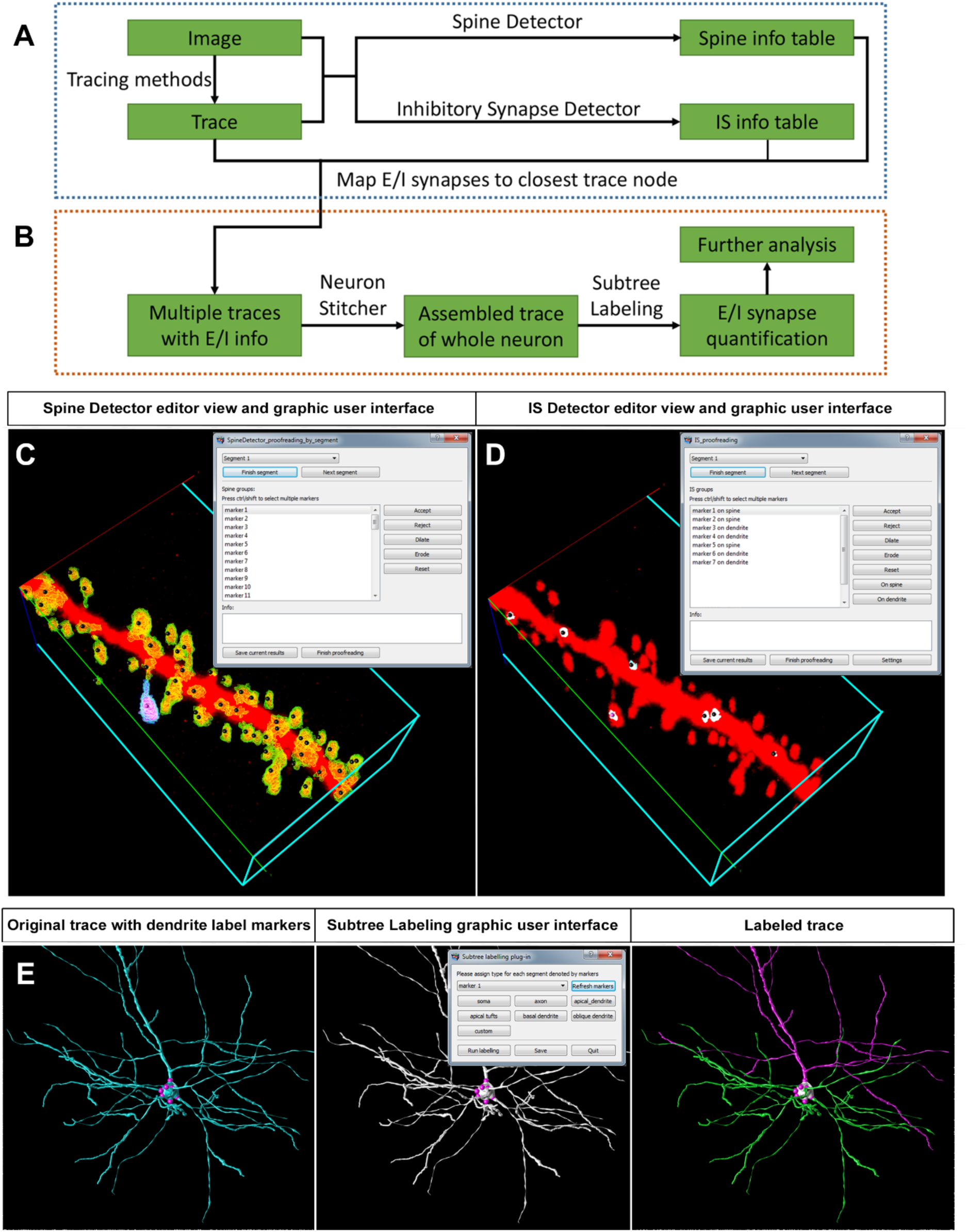
Synapse Detector computational pipeline and interactive user interface in Vaa3D. Related to Figure 1. **(A)** Extracting synaptic information from a neuron fragment within a single tissue section. **(B)** Mapping synaptic morphology across multiple tissue sections for whole-neuron spatial distribution analysis. **(C)** Spine Detector 3D annotation window and graphic user interface. Scale bar: 1 micron. **(D)** IS Detector 3D annotation window and graphic user interface. **(E)** User-directed annotation of dendritic domains with Subtree Labeling.

**Figure S2.**
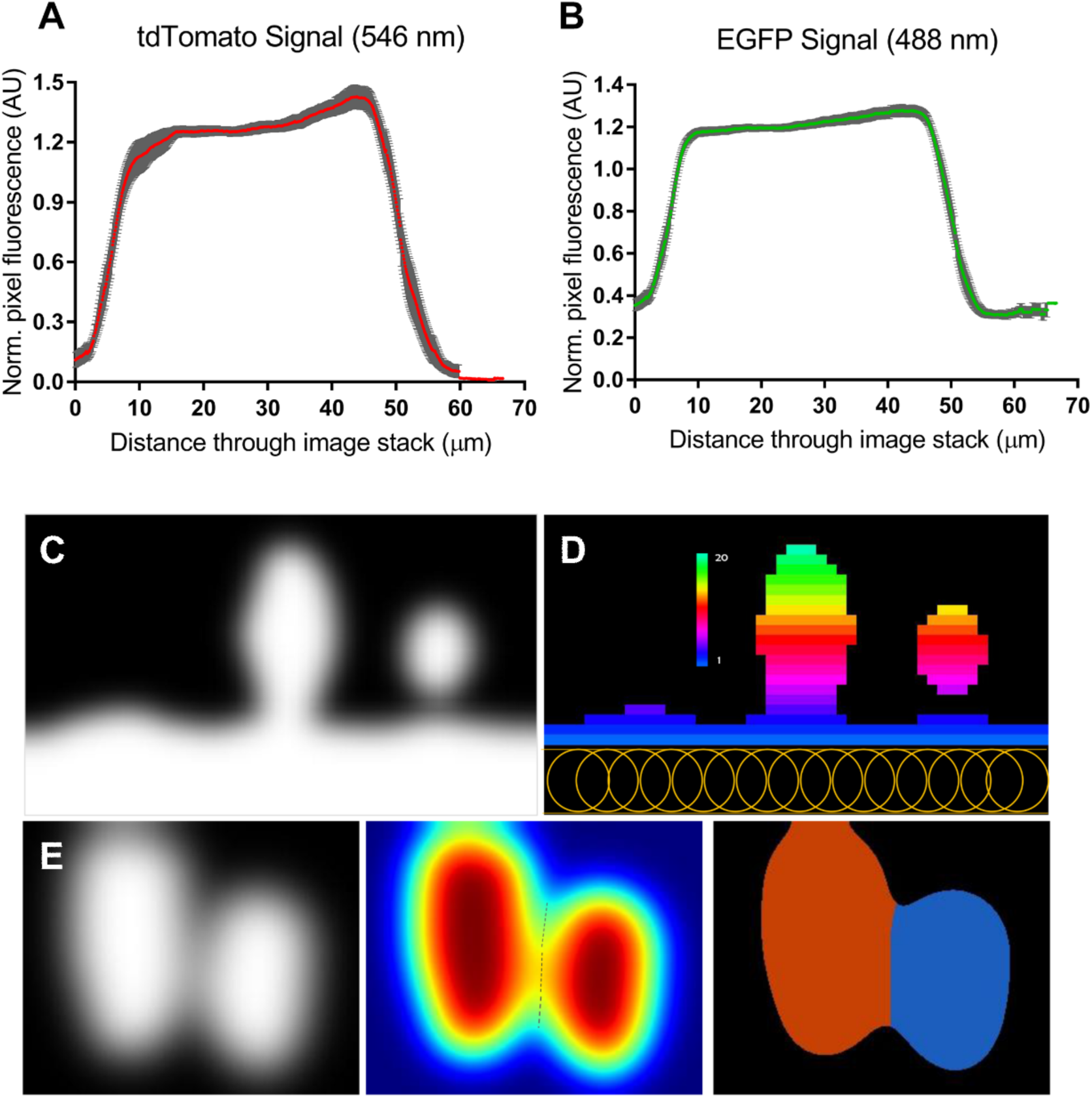
Synapse detection with confocal fluorescence imaging. Related to Figure 1. **(A)** Mean pixel fluorescence intensity through neuron fragment image stacks with 546 nm excitation wavelength. Distance normalized for tissue thickness. **(B)** Mean pixel fluorescence intensity through neuron fragment image stacks with 546 nm excitation wavelength. Distance normalized for tissue thickness. **(C)** Image fragment projected to 2D. **(D)** Color-coded distance to surface following distance transformation illustrates distance-based voxel clustering. Yellow circles represent the dendrite-filling neuron trace. **(E)** Intensity-based touching spine separation: 2D projection of two touching spines (left), intensity profiles of the two touching spines (middle), and separation of two touching spines (right).

**Figure S3.**
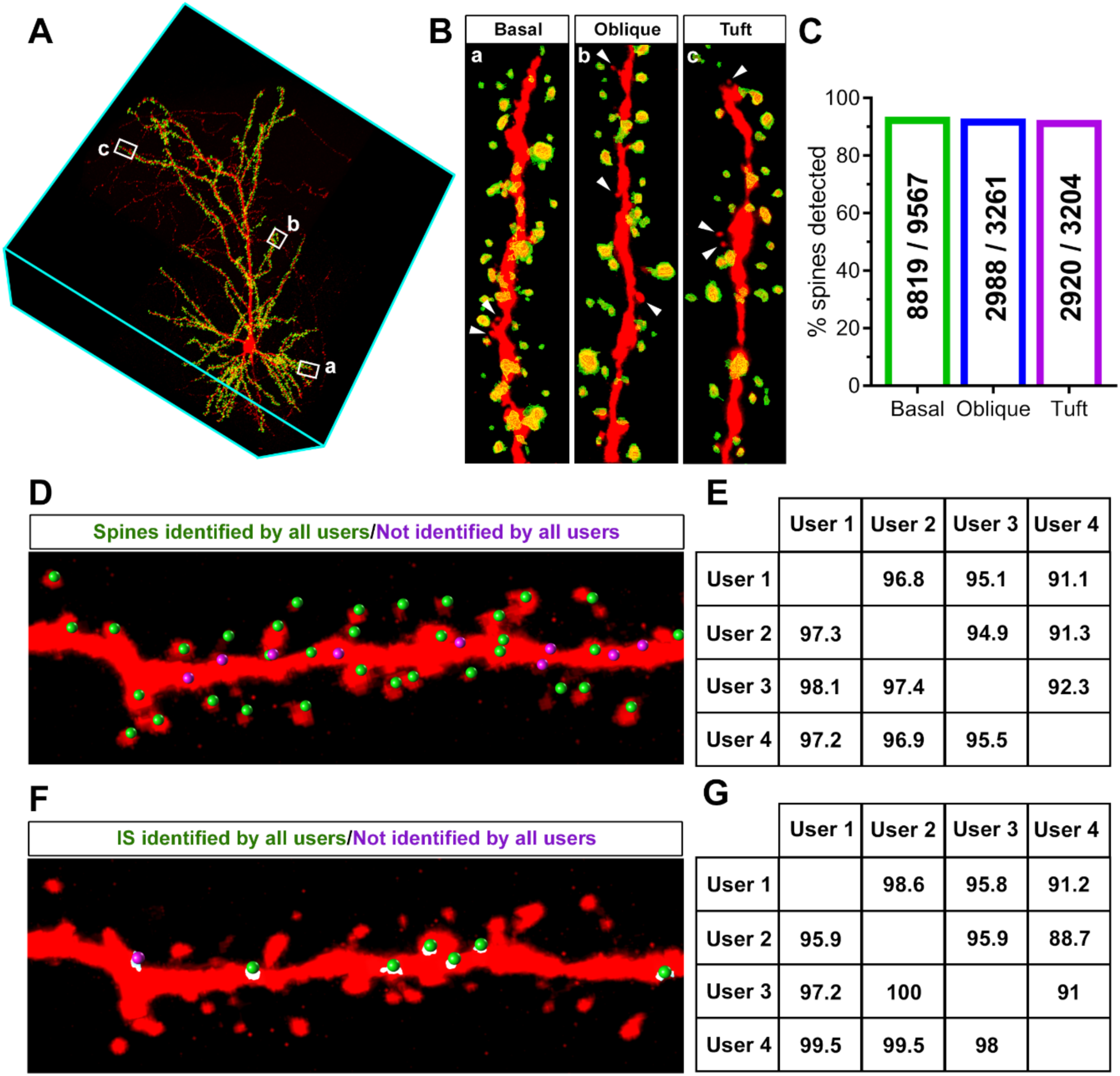
Synapse Detector false negative rate and results among multiple users. Related to Figure 1. **(A)** L2/3 PN reconstruction. **(B)** Examples of segments from dendrite domains in A. **(C)** Graph of spine detection rates for spines across dendritic domains. **(D)** Example segment with spines reconstructed by 4 independent experimenters. **(E)** Matrix showing percent agreement among experimenters reconstructing spines from basal dendrites. **(F)** Example segment with inhibitory synapses reconstructed by 4 independent experimenters. **(G)** Matrix showing percent agreement among experimenters reconstructing inhibitory synapses from basal dendrites.

**Figure S4.**
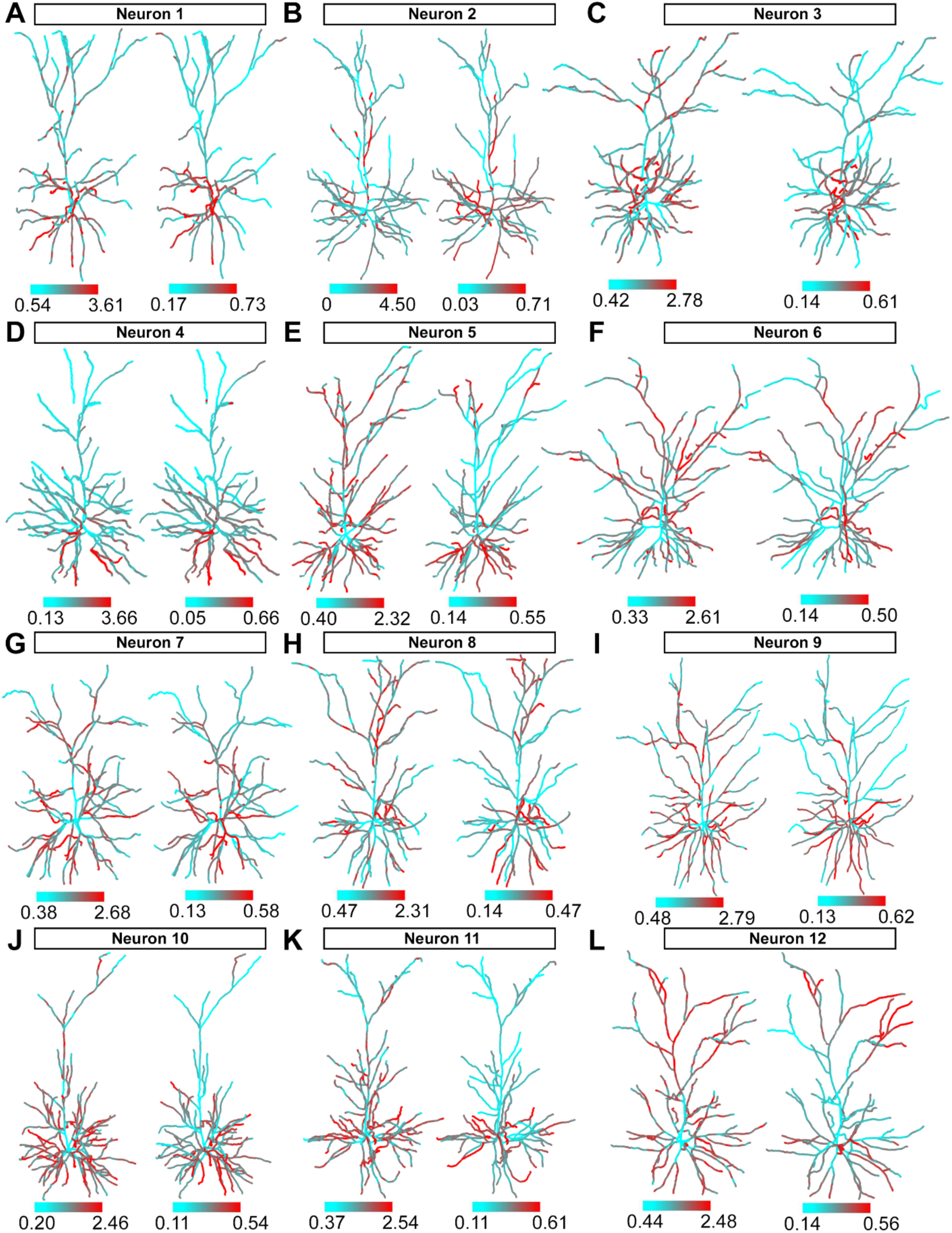
Neuron heat maps of excitatory synaptic density and large excitatory synaptic density. Related to Figure 3. **(A-L)** Left: heat maps of spine distribution indicating regions of low density in cyan and high density in red. Right: heat maps of large spine distribution. Large spines are defined as spines within the 20^th^ percentile for volume for each neuron individually. Density ranges are calculated for each neuron.

**Figure S5.**
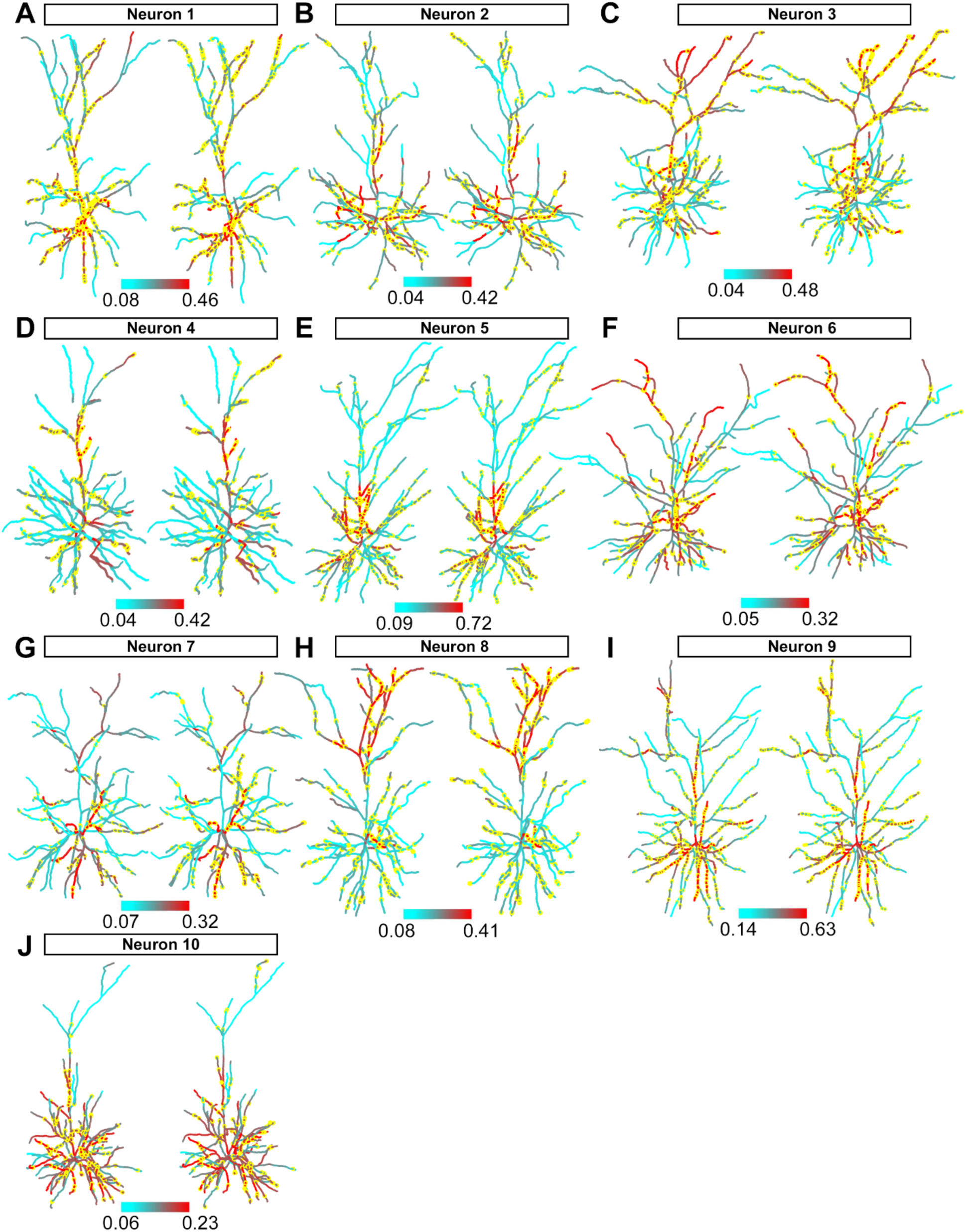
Neuron heat maps of large inhibitory synaptic distribution and inhibitory synaptic distribution on dendritic spines. Related to Figure 3. **(A-J)** Left: heat maps of inhibitory synaptic distribution where yellow puncta represent large inhibitory synapses. Large inhibitory synapses are defined as synapses within the 20th percentile for volume for each neuron individually. Right: heat maps of inhibitory synaptic distribution in which yellow puncta represent inhibitory synapses targeted to dendritic spines. Note the increased density of these dually-innervated spines toward the distal apical tufts and basal dendrites corresponding to thalamic input patterns. Density ranges are calculated for each neuron.

**Fig. S6.**
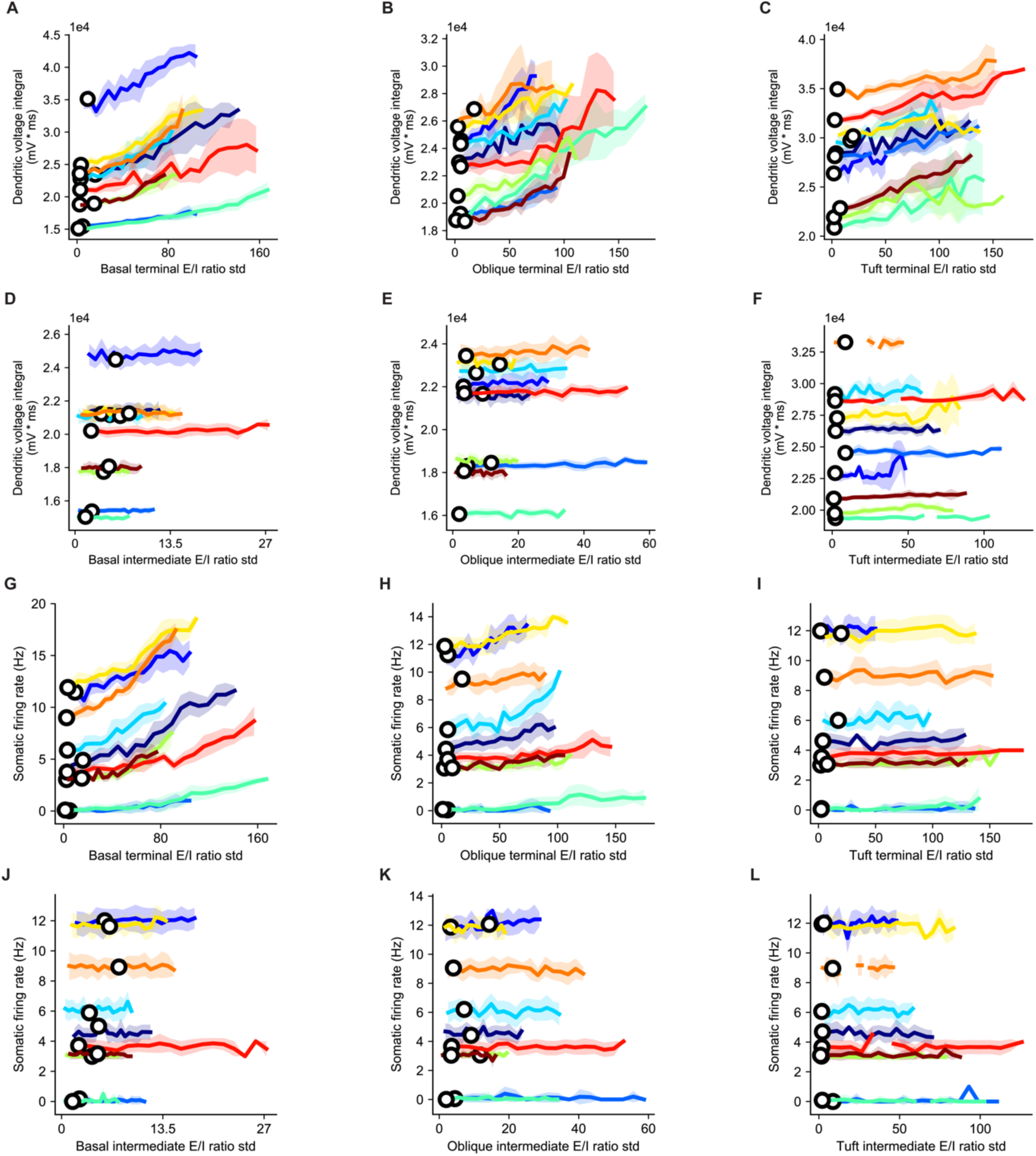
Terminal dendritic domains have balanced E/I ratio and are sensitive to variance in the E/I ratio among their respective dendritic branches. Intermediate dendritic domains have unbalanced E/I ratio and are not sensitive to the variance in their branch-specific E/I ratio. Related to Figure 7. **(A)** The average voltage time-integral for all branches in the basal terminal domain as a function of the standard deviation, SD, of the E/I ratio in this domain. Mean and SD are computed over ten different initial distributions of synapses (see **Materials and Methods**). The white circles represent the biological E/I SD. **(B-C)** Same as in **A,** with oblique terminal and tuft terminal domains. **(D)** As in **A**, for the basal intermediate domain. Note the variability in the biological E/I ratio variances and the insensitivity of the voltage time-integral for changes in E/I variance. **(E-F)** As in **D** for the oblique intermediate and tuft intermediate domains. **(G)** As is **A**, measuring the average somatic firing rate. **(H-I)** As in **G** for the oblique terminal and tuft terminal domains. **(J)** As in **G**, for the basal intermediate domain. **(K-L)** As in **J**, for the oblique intermediate and tuft intermediate domains.

**Fig. S7.**
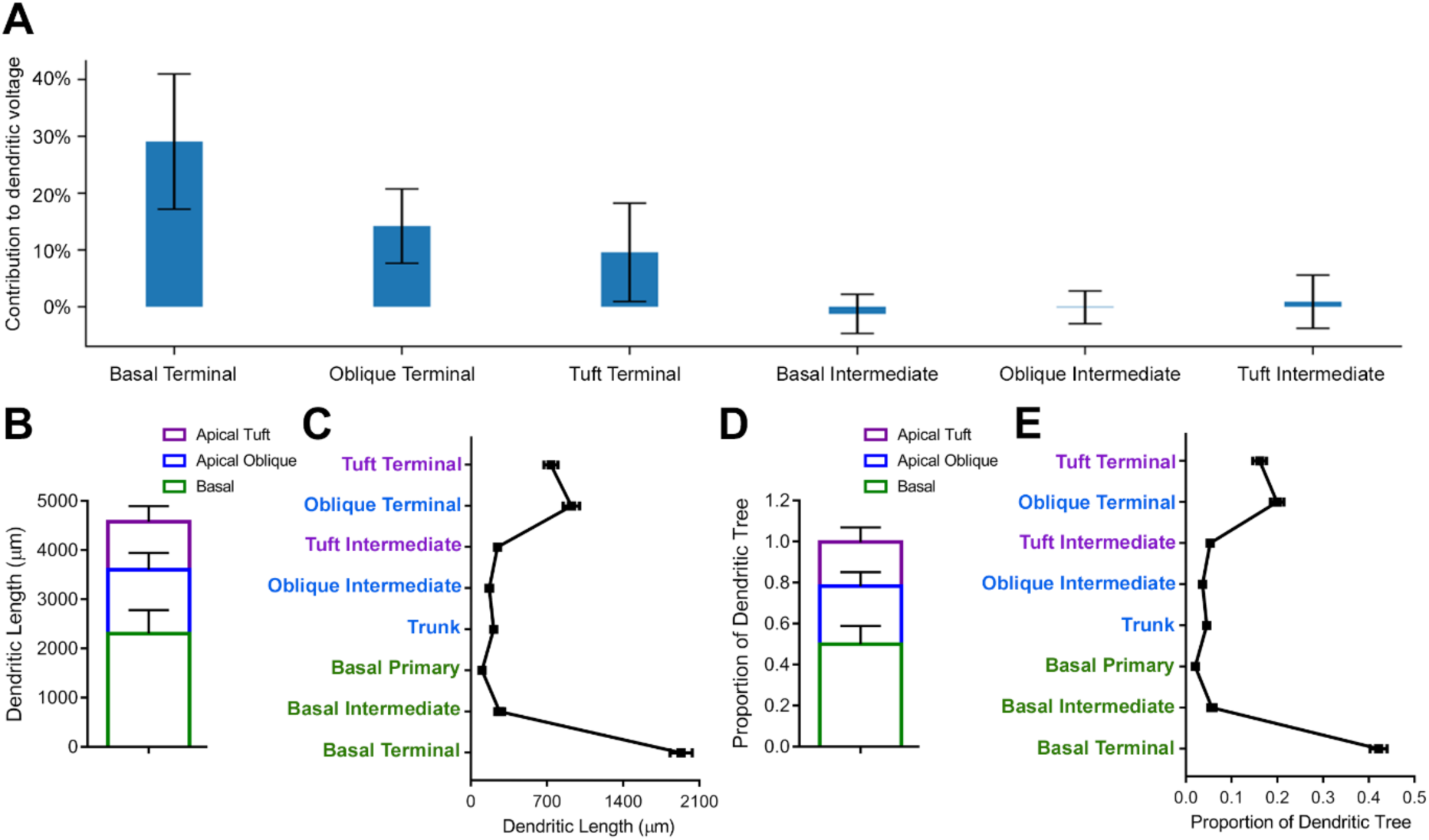
Branch type contribution to dendritic voltage and proportion of dendritic tree. Related to Figure 7. **(A)** The contribution of synapses located in a specific domain to the voltage time-integral in that domain (see **Materials and Methods**). Bars show the mean contribution over the ten neurons reconstructed and modeled, error bars show standard deviation of the contribution. **(B)** The total length of dendrites within domains across the dendritic tree. **(C)** The length of dendrites across dendritic branch types. **(D)** The proportion of the dendritic tree represented by each domain. **(E)** The proportion of the dendritic tree represented by each dendritic branch type. Note that these values correspond approximately to the contribution of synapses within each branch type to dendritic voltage.

## METHODS

Further information and requests for resources and reagents should be directed to and will be fulfilled by the Lead Contact, Franck Polleux.

## DATA ACQUISITION

### Mice

All animals were handled according to protocols approved by the Institutional Animal Care and Use Committee at Columbia University, New York. Postnatal day 42 CD-1 IGS mice (strain code: 022; Charles River) were used for all experiments. Timed-pregnant female mice were maintained in a 12 hour light/dark cycle and obtained by overnight breeding with males of the same strain. For timed-pregnant mating, noon after mating is considered E0.5.

### Constructs

The tdTomato reporter insert was subcloned into the pAAV-Ef1a-DIO eNpHR 3.0-EYFP plasmid (Addgene plasmid # 26966) between the AscI and NheI cloning sites. EGFP-GPHN (clone P1) was obtained from

H. Cline (TSRI, La Jolla, USA) and subcloned into pCAG downstream of a CMV-enhancer/chicken-β-actin (CAG) promoter, by replacing EGFP between the XmaI and NotI cloning sites.

### *In utero* electroporation

*In utero* cortical electroporation was performed at E15.5 on timed pregnant CD1 females. The previously described protocol for in utero cortical electroporation (*61*) was modified as follows. Endotoxin-free DNA was injected using a glass pipette into one ventricle of the mouse embryos. The volume of injected DNA was adjusted depending on the experiments. Electroporation was performed at E15.5 using a square wave electroporator (ECM 830, BTX) and gold paddles. The electroporation settings were: 5 pulses of 45 V for 50 ms with 500 ms intervals. Plasmids were used at the following concentrations: Flex-tdTomato reporter plasmid: 1 µg/µl; EGFP-GPHN 0.5 µg/µl; NLS-Cre recombinase: 0.0002 µg/µl.

### Tissue preparation

Animals at the indicated age were anaesthetized with isofluorane before intracardiac perfusion with PBS and 4% PFA (Electron Microscopy Sciences). 130 μm coronal brain sections were obtained using a vibrating microtome (Leica VT1200S). Sections were mounted on slides and briefly dehydrated at room temperature to reduce section thickness before being coverslipped in Fluoromount-G (SouthernBiotech).

### Confocal imaging

Confocal images of electroporated neurons in slices were acquired in 1024×1024 mode using an A1R laser scanning 11 confocal microscope controlled by the Nikon software NIS-Elements (Nikon Corporation, Melville, NY). We used a 100X H-TIRF, NA 1.49 (Nikon) objective lens to acquire image volumes of neuron fragments. Z-stacks of images were acquired with spacing of 100 nm. To counteract possible interference from light diffraction through the tissue, laser power was linearly increased as a function of depth within each tissue section to normalize the mean fluorescent intensity of pixels from image planes throughout the stack (**Figure S2A**). Dendritic spines and inhibitory synapses were quantified based on tdTomato fluorescence and EGFP-GPHN puncta fluorescence respectively. All quantifications were performed in L2/3 somatosensory cortex in sections of comparable rostro-caudal position.

### Heat map generation

We query the synaptic annotations for individual neurons to return a subset of the synapses satisfying the query. Some examples are “all inhibitory synapses,” “large spines,” and “all inhibitory synapses on spines.” We classified “large” synapses as greater than the 20^th^ percentile of synaptic volume for each neuron, closely corresponding to the persistent 160% increase in volume reported for synapses following structural forms of long-term potentiation (*5, 45, 46*). We calculate the path distances between these nodes and the soma, which is the 3D distance on the dendritic arbor from the soma to the node of interest. We calculate the distances between consecutive nodes that are in “ancestor-descendent” relationships on the neuronal arbor by obtaining the absolute value of the difference between their path distances.

To calculate the density of the synapses of interest at any given point on the dendritic arbor (the heat map), we count the synapses of interest that are within W µm of that point in terms of the path distance on the dendritic arbor, and convert these counts into color codes. Smaller W values increase the resolution of the heat map. On the other hand, when W is too small, the heat map will display high frequency noise. Therefore, we set the W values adaptively for each dendritic arbor as

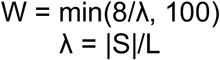

where |S| denotes the number of synapses of interest, and L denotes the total dendritic length of the arbor so that the unit of λ is μm^-1^. When mapping local counts to colors in the heat maps, we typically saturate the range of counts between the 2^nd^ and 98^th^ percentiles of the values to utilize the dynamic range of the colors more effectively.

### E/I balance heat map generation

The excitatory and inhibitory heat map values for individual neurons are scaled and shifted to lie in the *[0, 1]* interval. The absolute value of the difference, which again lies in the *[0, 1]* interval, is displayed.

### Within-domain randomization for structural organization analysis

For each neuron, we first find all the nodes of the arbor trace in the domain of interest. The nodes that carry synapses on them have extra annotations reflecting the size and type of the synapses. Then, we reassign the size-and-type annotations to those nodes uniformly at random, thus leaving the structure of the arbor unchanged.

### Branch-level synaptic rate correlation analysis

For each relevant branch (i.e., primary, intermediate, terminal) in each neuron, we count the synapses of interest and divide by the path length of that branch to obtain the density estimate. We calculate the correlation coefficient and the *p*-value pair for each plot using the *corrcoef* command in MATLAB.

### Analysis software

The software used to generate the heat maps, the rate plots, and the within-domain randomization results is available at https://github.com/uygarsumbul/spines.

### Quantification and statistical analysis of synaptic distribution data

Data is shown as the mean ± SEM, unless otherwise stated. T-tests was used to compare the mean of two groups with corrections for multiple comparisons: discovery determined using the Two-stage linear step-up procedure of Benjamini, Krieger and Yekutieli, with Q = 1%. A one-way ANOVA was used when more than two groups existed. Significance for all experiments was placed at p < 0.05. Statistical tests were carried out with GraphPad Prism.

### Modeling

Reconstructed morphological data, synaptic attributes and spatial distribution of E and I synapses were taken from Vaa3D reconstructions. Modeling and simulation was performed using NEURON simulator, accessed using a python script (*62*). Specific membrane resistance and capacitance, and axial resistance were 12,000 Ωcm^2^, 1 µF/cm^2^, 150 Ωcm, respectively. These values were chosen such that the somatic input resistance and time constant will be within known biological ranges for these neurons (92 ± 15*M*Ω and 12 *ms*, respectively (*63*)). Active membrane ion channels were taken from the Blue Brain Project models of L2/3 PNs (*64*) and tuned to produce similar results to that found *in vivo* for L2/3 PNs (*60*). The activation of excitatory and inhibitory synapses was randomly sampled from a Poisson distribution with an average of 1.75 Hz and 10 Hz for the E and I synapses, respectively. This generated a mean somatic firing rate for the 10 modeled cells of 4.6 ± 3.6 Hz, similar to that found experimentally (*31*). The synaptic peak conductance for the Esynapses was 0.4 nS (for α-amino-3-hydroxy-5-methyl-4-isoxazolepropionic acid (AMPA) component as well as for the N-methyl-D-aspartate (NMDA) components) and 1 nS for the γ-Aminobutyric acid (GABAA) synapses. The rise time constants for the conductances of these synapses was 0.2 ms, 2.04 ms and 0.18 ms, respectively and the respective decay time was 1.7 ms, 75.2 ms and 1.7 ms. The reversal potential values are 0 mV, 0 mV, and −80 mV, respectively. Dendritic voltage traces were recorded from the center of the respective dendritic branch.

### Fitting model to *in vitro* experiment results

To ensure that our model captures important aspects of dendritic nonlinearities and voltage attenuation, we tuned the *Na+* and *Ih* membrane conductances to replicate two experiments as in Waters et al, 2003. In order to replicate the attenuation of the back-propagating action potential along the apical trunk in L2/3 PN as in **Figure 6C**, a step depolarization current of 200 pA for 200 ms was injected to the modelled soma, invoking a somatic action potential, and recorded the amplitude at 10 µm intervals along the apical trunk (**Figure 6D**). To replicate the contribution of Na^+^ channels to the backpropagation of action potentials (**Figure 6E**), we have injected 200 pA for 200 ms to the soma, invoking an action potential, and recorded the voltage both at the soma and 80 µm from the soma on the apical trunk. Then, we simulated the application of TTX by removing the Na^+^ channels from the model, voltage clamping the soma to the voltage trace created by the action potential, and recording the amplitude of the passively propagated action potential 80 µm from the soma on the apical trunk (**Figure 6F**). The model (**Figures 6D** and **6F**) was able to replicate the experimental results (**Figures 6C** and **6E**) of both the attenuation and the dependency on Na+ channels of the back-propagating action potential.

As found experimentally (*31*), our L23 PN models produced a range of firing rates; some cells fire at high rate (10 Hz, red circle in **Figures 6G-6I** recorded) and some fire at low rate (3 Hz, blue circle in **Figures 6G-6I**). This variance in somatic firing rate persisted despite that fact that all models have the same passive and active properties and E and I input frequencies. We found that the global E/I balance per cell (**Figure 6G**) was a strong indicator of the output firing rate (**Figure 7F**). Additionally, we found that the somatic firing rate is correlated with the size of the cell, the larger the surface area of the cell, the lower is its firing rate (**Figure 6H**). This is due to the interesting experimental finding that larger cells have lower global E/I ratio and, consequently, that their firing rate is lower (**Figure 6I**).

### Changing E/I ratio variance across dendritic domains

To study the influence of the ratio of excitatory and inhibitory synapses in a given dendritic branch, we iteratively increased or decreased the variance of E/I ratios over all branches belonging to a given domain. To change the E/I ratio variance, we randomly distributed the location of synapses between branches, while keeping the total number of synapses in the domain fixed (as found experimentally for the respective modeled cell). This process was repeated ten times, each time with a different initial distribution of the synapses. In **Figures 7D** and **7E** and **Figure S6** the voltage time integral (in a time window of 3000 sec) was computed at the center of each branch in a particular domain, for different E/I SD, averaged over all branches in that domain. The same was performed for the somatic firing rates (**Figure 7F**) for different E/I SD in the basal terminal domain.

### Contribution of synapses in a domain to voltage in that domain

To measure the contribution of synapses located in a specific domain to the depolarization in that domain, we simulated each of the modeled cell with excitatory and inhibitory synapses as described above, and calculated the mean voltage time-integral in each domain. We then calculated the respective mean voltage time-integral when all synapses in that domain were not active and compared the two cases (**Figure S7A**).

## SOFTWARE DEVELOPMENT

### Computational pipeline overview

Information for dendritic spine placement and morphology was acquired from large-volume high-resolution image stacks of thick brain tissue. There are two possible strategies for quantifying the spatial distribution of excitatory and inhibitory (E and I) synapses of an entire neuron:

1. Stitch image volumes together prior to analysis.
2. Analyze each image volume independently and align the spatial information recorded from each image to create a complete neuron representation.

The first strategy, which involves all the image stacks into a terabyte volume and then perform neuron tracing, synapse segmentation and spatial analysis globally on the combined volume. The downside of the approach is a big data problem of manipulating, storing and analyzing the giant volume. Additionally, this approach is computationally wasteful because only a fraction of the stitched volume contains relevant structure. To avoid this big data problem, we pursued the alternative strategy of performing dendrite tracing and synapse segmentation on each image stack individually and associating morphological information of each synapse to a specific node of the trace (thereby encoding the location of every synapse within the spatial context of the neuron). To create representations of complete neurons across serial vibratome sections, dendrite traces containing synaptic information were aligned and stitched together. In comparison with the terabyte combined volume generated by the first reconstruction strategy, the resulting reconstructions are 4-6 megabytes in size.

Our pipeline for whole-neuron synaptic reconstruction consists of two parts. In the first part, we extract E and I synaptic information for an individual image across a tissue section (**Figure S1A**). For each image stack, we trace the dendritic arbor of the neuron fragment using automatic tracing methods followed by manual corrections. Then both Spine Detector and IS Detector take the neuron skeleton and the image stack as input to automatically isolate spines and inhibitory synapses within a user-defined radius of each dendrite. Spine Detector generates a table that records the local information of dendritic spines including the distance between each synapse and the dendrite, volume, and the nearest tree node. IS Detector generates a table that records the local information of inhibitory synapses including volume, whether the inhibitory synapse is located on a spine or the dendrite, and the nearest tree node. These morphological characteristics of synapses impact their neurotransmitter content and integration properties (*43, 65-67*).

In the second part of our reconstruction pipeline, we map E and I synaptic morphology across multiple images for whole-neuron spatial distribution analysis (**Figure S1B**). First, the dendritic spine information and inhibitory synaptic information from each image are mapped to the closest tree node of their corresponding dendrite trace. Next, the traces containing local synaptic information from each image stack are aligned and stitched together to generate a whole-neuron synaptic reconstruction. Notably, the association between synapses and their respective tree nodes remains unchanged during the assembly. After obtaining the single reconstruction trace of the whole neuron, we subtype the dendritic arbor in terms of identity and morphology so that we can analyze the synaptic features within domain and segment levels.

### Neuron reconstruction

Digital reconstructions, or traces, are an effective representation of neuronal topology and geometry. The traces are usually described using a tree graph and consist of 3-D point coordinates, diameters, and connectivity between points. This succinct representation enables an extensive quantitative analyses of the geometrical organization of the neurons they represent including total length, branching angles, distribution statistics and cumulative distance from the soma (*68*). Numerous automated tracing methods have been developed (*69-71*). In this paper, the initial reconstructions are obtained using the automatic tracing methods built in the open source 3D visualization and analysis tool Vaa3D (*52*). Then, experts manually proofread the traces and make adjustments with the built-in proof-editing tools. Notably our synapse analysis pipeline works for traces generated by all tracing methods. Accurate reconstructions are important to improve the performance of automatic synapse detection.

### Automatic spine detection

To automatically identify potential spines, Spine Detector segments candidate spine-associated voxels whose fluorescence is greater than a linearly interpolated local threshold between nodes along the closest dendritic segment (*72*). Spine detection is performed within a user-defined region around the dendrite and intensity threshold such that all voxels within the user-defined region and above the threshold are identified possible spine voxels. Spine Detector takes both the image and the dendritic trace as input and clusters adjacent voxels in the cell-fill channel based on their distance from the dendrite surface. Touching spines are separated based on voxel intensities. Because the dendrite traces represent the dendrites with a series of overlapping nodes (*73*), information about the volume and distance from the dendrite of each spine can be associated with its nearest node to assign a location within the spatial context of the dendritic arbor.

### Voxel clustering for enhanced detection

In contrast to previous approaches that estimate spine volume from the spine tip backward toward the dendrite (*72*), Spine Detector identifies potential spine voxels at the dendrite shaft and estimates their volume by iteratively adding layers of connected voxels toward the spine tips. To quickly estimate the minimum distance between each voxel and the nearest dendrite surface, Spine Detector uses the radius of each node across the neuron trace as a representation of the dendrite surface and performs a distance transform on the image (**Figures S2C** and **S2D**). The initial seeds of potential spines are the voxels the shortest distance from the dendrite surface. In each iteration, potential spines are identified and grown by adding new layers of connected neighbor-voxels until a spine edge is detected. This is achieved by establishing a floor value to the distance between the initial seeds and the dendrite surface and repeatedly adding layers of connected neighbor-voxels equal to the floor value of the previous layer. At the end of each iteration, Spine Detector determines whether the number of voxels have exceeded the user-defined spine size and whether the maximum layer width has exceeded the user-defined layer width. If the most recently added layer did not meet these criteria, all previous layers are discarded and the voxels in that layer serve as the seed for the next layer. The iteration stops when all qualified voxels are assessed. Spine candidates are rejected based on user-supplied parameters for minimum voxel count and minimum spine length, allowing users to reconstruct images acquired at different magnifications. Notably, spines can be detected with this methodology regardless of the resolution of the spine neck.

### Intensity-based segmentation of adjacent spines

Limited image resolution, inaccurate thresholding, and physical proximity can all give rise to adjacent spines incorrectly categorized as a single synapse. Based on the observation that spine voxel intensities are naturally brighter at the center than the edges, we adopted an adapted watershed algorithm to separate spines within close spatial proximity (*74*). First an initial threshold is set at a relatively high fluorescence intensity so that only the center-voxels of spines are identified (**Figure S2E**). With the successively decreasing fluorescence toward the spine border, the spine boundary grows in size. When two potential spine boundaries meet they each become defined to separate adjacent spines. The merger of two spine volumes is only considered when both spines are relatively small (lower than 1% of the average volume).

### Inhibitory synapse detection

We labeled inhibitory synapses using the scaffolding protein Gephyrin tagged with a fluorescent protein as a marker (*9*). Because these synapses can only occur on the dendrites or the spines of neurons of interest, we use the image from the cell-fill channel containing the dendrites and the spines as a mask image to extract the relevant region for the inhibitory synaptic marker. Then, signal beyond user-input parameters for minimum/maximum voxel count and distance from the trace is excluded and potential inhibitory synapses from the resulting image are identified based on a user-input intensity threshold. Users have the ability to accept or reject potential inhibitory synapses, adjust their volume, and assign them as dendrite-targeting or spine-targeting.

### Stitching neuron traces across serial 3D image sections

To assemble the neuron reconstructions traced across multiple image stacks we used Neuron Stitcher (*36*), a software suite for stitching non-overlapping neuron fragments in serial 3D image sections. The software identifies severed neurite traces at the section planes, known as ‘border tips’, and then uses a triangle matching algorithm to align traces created from neurons spanning serial tissue sections. Once the initial border tip matches are identified, the alignment is estimated in the form of an affine transformation and the border tips are connected to form a complete neuron trace.

### Neurite subtyping

To better understand the synaptic distribution within domain and segment levels, we developed the Subtree Labeling program as a plug-in of Vaa3D to subtype neurites for further analysis. Using this program it is possible to assign a neurite segment into multiple categories: axon, soma, apical trunk, apical tufts, apical oblique dendrites, and basal dendrites. The user interface allows the user to select the starting vertex for each branch and to assign neurite type. The program first finds the tree node for soma and sorts the tree with the soma node as the tree root. Then, all the child vertices of the starting vertex are assigned the same branch type as each manually annotated starting vertex.

## USER’S GUIDE

### Synapse Detector Interactive User Interface

To broaden the utility of SynapseDetector to work with a variety of different data acquisition processes, we designed an interactive interface to (1) allow visual evaluation of detection results and accept or reject putative synapses; and (2) enable manual correction of synaptic volume through addition or subtraction of associated pixels. The software was implemented in C/C++ as a plugin of Vaa3D, which is a publicly available open source platform with a user-friendly interface for 3D+ image analysis and visualization. In the following sections, we will introduce how to use the tools. For detailed directions how to create neuron traces using Vaa3D, see the recently published protocol (*52*).

Main website: http://vaa3d.org/

Documentation: http://code.google.com/p/vaa3d/

Help/DiscussionForum: http://www.nitrc.org/forum/forum.php?forum_id=1553

Bug tracking and requesting new features: http://www.nitrc.org/tracker/?group_id=379

### Sorting dendrite traces for reconstruction

A dendrite trace (swc file) is composed of a series of connected nodes with varying radii. This plugin connects nodes that were not linked during manual trace editing, which is critical for proper segment classification. This plugin allows the user to designate the soma as the “root node,” the first node in the tree from which the distance to all daughter nodes can be determined to analyze synaptic distribution.

1. In Vaa3D, drag a neuron trace into the 3D viewer.
2. Use ‘Cmd/Ctrl+L’ to toggle between the line (skeleton) display mode and the surface mesh display mode of the neuron. In line display mode it is possible to visualize root nodes contained within the trace.
3. If the trace contains a soma, hover cursor over soma to identify the node number that will be designated as the root node.
4. In Vaa3D, go to the ‘Plug-in’ main window menu and click ‘neuron_utilities’, then click on ‘sort_neuron_swc’, and finally click on ‘sort_swc’.
5. Select the trace in the ‘Open from 3D Viewer’ tab.
6. If the trace contains a soma, specify the root node number as the soma node number. If the trace does not contain a soma, click ‘cancel’.
7. Specify a voxel threshold for adjacent segments to be connected. To connect all segments click ‘cancel’. Save the sorted neuron trace.

### Resampling dendrite traces for reconstruction

To maximize the spatial resolution of synaptic distribution analysis, it is recommended to resample the associated neuron trace to contain the highest possible number of tree nodes.

1. In Vaa3D, go to the ‘Plug-in’ main window menu and click ‘neuron_utilities’, then click on ‘resample_swc’, and finally click on ‘resample’.
2. Select the trace and specify a step length of 1. Click ‘ok’ and save the resampled neuron trace.

### Using Crop Image Trace to analyze large image volumes

This new tool allows the user to analyze image volumes with Synapse Detector that would normally be too large by cropping a region of interest based on XYZ pixel coordinates and aligning an associated neuron trace to the resulting image volume. In practice, image volumes greater than 2000 x 2000 pixels in X and Y and 500 pixels in Z are difficult to reconstruct without cropping.

1. In Vaa3D, use ‘Cmd/Ctrl+O’ to open the appropriate image file.
2. In the tri-view window, click ‘see in 3D’ and then click ‘entire image’ to visualize the image file.
3. Drag and drop the neuron trace corresponding to the image file into the 3D view window.
4. Go to the ‘Plug-in’ main window menu and click ‘image_geometry’,andthen‘crop_image_tace’, and finally click on ‘crop’.
5. Select an appropriate output directory, and specify the XYZ coordinates to crop the image (the number of pixels in X, Y, and Z that compose each image can be viewed in the tri-view window and 3D viewer), and specify the color channels to include in the new image. Click on ‘run and save’.

### Synapse annotation with Synapse Detector

This new tool semi-automatedly identifies dendritic spines (Spine Detector) or inhibitory synapses (IS Detector) and quantifies their morphology and spatial distribution. Synapses can be manually accepted or rejected, as well as edited by dilating or eroding pixels. IS Detector also allows the user to designate inhibitory synapse location on either a spine or the dendritic shaft.

### Spine Detector user interface

1. In Vaa3D, go to the ‘Plug-in’ main window menu, click ‘synapse_detector’, and click on ‘SpineDetector_NewProject’. Users can also continue an existing project by clicking ‘SpineDetector_ExisitingProject’.
2. Load the image volume (v3dpbd or v3draw), associated trace file (swc), and designate an output destination for the sorted reconstruction. Select the color channel of the cell-fill.
3. Specify the threshold for background signal in the image and volume parameters for potential spines. Pixel to micron conversion can be calculated from the imaging magnification and is usually stored within the image properties. Click ‘Run’.
4. Click ‘Proofread by segment’ to edit spines along a dendrite segment (recommended) or ‘Proofread by spine’ to edit each spine individually.
5. Accept/reject potential spines and proofread spine morphology by dilating/eroding volume. The highlighted regions indicate the potential spines (**Figure S1C**). It is recommended to look at the segment at different angles and toggle between views with the spine annotation channel on and off in the 3D viewer.
6. Click ‘Save current result’ to save intermediate results during proofreading. Spine Detector will generate 4 files in the output folder: a text file ‘project.txt’ (includes all info needed to reload the last saved reconstruction project), a marker file indicating the positions of accepted/rejected spines, a csv file (table of accepted spine information), and an image file of accepted spines.
7. Click ‘Finish proofreading’ to save final results after proofreading. After proofreading is completed, Spine Detector generates 2 image files (edited spine reconstruction and isolated spine annotations), a marker file of spine positions, and a csv file containing spine morphology data (all data measured in pixels).

### IS Detector user interface

1. In Vaa3D, go to the ‘Plug-in’ main window menu, click ‘synapse_detector’, and click on ‘IS_Detector_NewProject’. Users can also click on ‘IS_Detector_ExisitingProject’ to reload a previously saved project.
2. Load the image volume (v3dpbd or v3draw), associated trace file (swc), and designate an output destination for the sorted reconstruction. Select the color channel of the cell-fill and the color channel of the inhibitory synaptic marker (or other punctate intracellular marker).
3. Specify the threshold for background signal in both image channels and volume parameters for potential inhibitory synapses. Pixel to micron conversion can be calculated from the imaging magnification and is usually stored within the image properties. Click ‘Run’, and then click ‘Proofread by segment’.
4. Accept/reject potential inhibitory synapses and proofread morphology by dilating/eroding volume and specifying synapse location on spine/dendrite. The highlighted regions indicate the potential inhibitory synapses (**Figure S1D**). It is recommended to adjust the lookup table thresholds for synaptic visualization by clicking the ‘Vol Colormap’ button on the right-side control pane of the 3D viewer.
5. Click ‘Save current result’ to save intermediate results during proofreading. Spine Detector will generate 2 files in the output folder: a text file (includes all info needed to reload the last saved reconstruction project) and a csv file (table of accepted spine information).
6. Click ‘Finish proofreading’ to save final results after proofreading. After proofreading is completed, Spine Detector generates 2 image files (unedited and edited inhibitory synapses), a marker file of synaptic positions, and a csv file containing synaptic morphology data (all data measured in pixels).

### Embedding synaptic data within the neuron trace

After spines and inhibitory synapses have been annotated throughout the image volume, synaptic information stored in tables can be associated with their corresponding nodes throughout the neuron trace using the Synapse Detector Combiner.

1. In Vaa3D, go to the ‘Plug-in’ main window menu click ‘synapse_detector’, and click on ‘Combiner’.
2. Load the spine and inhibitory synapse tables that correspond to the neuron trace. If the image volume was cropped before reconstruction the trace will be associated with tables from each cropped region. Click ‘Run’, and save the neuron reconstruction.

### Assembling reconstruction fragments with Neuron Stitcher

For detailed directions how to stitch neuron traces using Neuron Stitcher, see the recently published protocol (*36*). To preserve synaptic information within the reconstruction, assemble reconstruction fragments to produce an eswc trace.

### Annotating reconstruction traces with Subtree Labeling

This new tool creates an enhanced neuron skeleton that contains information about dendrite identity, branch order, and cumulative dendritic distance from the soma. The user interface allows the user to select the starting vertex for each branch and to assign neurite type. Child vertices of each starting vertex are assigned the same branch type as each manually annotated starting vertex. To label neuron reconstructions stitched from multiple fragments throughout the entire dendritic arbor, these traces must first be sorted with the Neuron Connector plugin to preserve the eswc file type.

1. In Vaa3D, go to the ‘Plug-in’ main window menu,click‘neuron_utilities’and‘neuron_connector’,andselect‘connect_neuron_swc’.
2. Load the input trace file and designate an output destination for the sorted reconstruction.
3. Set the ‘connection configuration’ to ‘connect all, shortest distance and click ‘Connect’.
4. After sorting the reconstruction, drag it into the 3D viewer.
5. Use ‘Cmd/Ctrl+L’ to toggle between the line (skeleton) display mode and the surface mesh display mode of the neuron. In line display mode it is possible to visualize root nodes contained within the trace.
6. Right-click at the soma to and click ‘create marker from the nearest neuron-node’ to create a marker at the root node. Create markers between the root node and dendrite terminals according to experiment-specific labeling schemes (**Figure S1E**).
7. In Vaa3D, go to the ‘Plug-in’ main window menu, click ‘neuron_utilities’, and then ‘subtree_labeling’.
8. Select ‘Refresh markers’ to ensure all markers were selected. Assign dendrite labels to each marker. It is possible to add new markers and click ‘Refresh markers’ to add them to the list of labeled markers. Markers will be labeled in descending order starting with marker 1, so it is recommended to place makers from the root node outward according to the labeling scheme.
9. Click ‘run labeling’. Review the neuron trace in the 3D viewer to verify segments were properly labeled and click ‘save’ within the Subtree Labeling interface window.

## REFERENCES

1. A. Polsky, B. W. Mel, J. Schiller, Computational subunits in thin dendrites of pyramidal cells. Nat Neurosci 7, 621–627(2004).

2. G. Liu, Local structural balance and functional interaction of excitatory and inhibitory synapses in hippocampal dendrites. Nat Neurosci 7, 373–379 (2004).

3. Y. Katz et al., Synapse distribution suggests a two-stage model of dendritic integration in CA1 pyramidal neurons. Neuron 63, 171–177 (2009).

4. A. Gidon, I. Segev, Principles governing the operation of synaptic inhibition in dendrites. Neuron 75, 330–341 (2012).

5. C. D. Harvey, K. Svoboda, Locally dynamic synaptic learning rules in pyramidal neuron dendrites. Nature 450, 1195–1200 (2007).

6. T. Kleindienst, J. Winnubst, C. Roth-Alpermann, T. Bonhoeffer, C. Lohmann, Activity-dependent clustering of functional synaptic inputs on developing hippocampal dendrites. Neuron 72, 1012–1024 (2011).

7. H. Makino, R. Malinow, Compartmentalized versus global synaptic plasticity on dendrites controlled by experience. Neuron 72, 1001–1011 (2011).

8. A. C. Frank et al., Hotspots of dendritic spine turnover facilitate clustered spine addition and learning and memory. Nat Commun 9, 422 (2018).

9. J. L. Chen et al., Clustered dynamics of inhibitory synapses and dendritic spines in the adult neocortex. Neuron 74, 361–373 (2012).

10. M. F. Iacaruso, I. T. Gasler, S. B. Hofer, Synaptic organization of visual space in primary visual cortex. Nature 547, 449–452 (2017).

11. Y. M. Sigal, C. M. Speer, H. P. Babcock, X. Zhuang, Mapping Synaptic Input Fields of Neurons with Super-Resolution Imaging. Cell 163, 493–505 (2015).

12. D. G. C. Hildebrand et al., Whole-brain serial-section electron microscopy in larval zebrafish. Nature 545, 345–349 (2017).

13. E. Meijering, A. E. Carpenter, H. Peng, F. A. Hamprecht, J. C. Olivo-Marin, Imagining the future of bioimage analysis. Nat Biotechnol 34, 1250–1255 (2016).

14. D. Kleinfeld et al., Large-scale automated histology in the pursuit of connectomes. J Neurosci 31, 16125–16138 (2011).

15. M. Helmstaedter, Cellular-resolution connectomics: challenges of dense neural circuit reconstruction. Nat Methods 10, 501–507 (2013).

16. M. J. Fogarty, L. A. Hammond, R. Kanjhan, M. C. Bellingham, P. G. Noakes, A method for the three-dimensional reconstruction of Neurobiotin-filled neurons and the location of their synaptic inputs. Front Neural Circuits 7, 153 (2013).

17. D. L. Dickstein et al., Automatic Dendritic Spine Quantification from Confocal Data with Neurolucida 360. Curr Protoc Neurosci 77, 1 27 21–21 27 21 (2016).

18. B. J. Marlin, M. Mitre, A. D’Amour J, M. V. Chao, R. C. Froemke, Oxytocin enables maternal behaviour by balancing cortical inhibition. Nature 520, 499–504 (2015).

19. R. C. Froemke et al., Long-term modification of cortical synapses improves sensory perception. Nat Neurosci 16, 79–88 (2013).

20. A. L. Dorrn, K. Yuan, A. J. Barker, C. E. Schreiner, R. C. Froemke, Developmental sensory experience balances cortical excitation and inhibition. Nature 465, 932–936 (2010).

21. A. D’Amour J, R. C. Froemke, Inhibitory and excitatory spike-timing-dependent plasticity in the auditory cortex. Neuron 86, 514–528 (2015).

22. M. J. Higley, D. Contreras, Balanced excitation and inhibition determine spike timing during frequency adaptation. J Neurosci 26, 448–457 (2006).

23. M. Xue, B. V. Atallah, M. Scanziani, Equalizing excitation-inhibition ratios across visual cortical neurons. Nature 511, 596–600 (2014).

24. N. Takahashi, C. Kobayashi, T. Ishikawa, Y. Ikegaya, Subcellular Imbalances in Synaptic Activity. Cell Rep 14, 1348–1354 (2016).

25. C. Q. Chiu et al., Input-Specific NMDAR-Dependent Potentiation of Dendritic GABAergic Inhibition. Neuron 97, 368–377 e363 (2018).

26. I. Ballesteros-Yanez, R. Benavides-Piccione, G. N. Elston, R. Yuste, J. DeFelipe, Density and morphology of dendritic spines in mouse neocortex. Neuroscience 138, 403–409 (2006).

27. Y. Kubota, S. Hatada, S. Kondo, F. Karube, Y. Kawaguchi, Neocortical inhibitory terminals innervate dendritic spines targeted by thalamocortical afferents. J Neurosci 27, 1139–1150 (2007).

28. L. A. DeNardo, D. S. Berns, K. DeLoach, L. Luo, Connectivity of mouse somatosensory and prefrontal cortex examined with trans-synaptic tracing. Nat Neurosci 18, 1687–1697 (2015).

29. L. Frangeul et al., A cross-modal genetic framework for the development and plasticity of sensory pathways. Nature 538, 96–98 (2016).

30. S. Lefort, C. Tomm, J. C. Floyd Sarria, C. C. Petersen, The excitatory neuronal network of the C2 barrel column in mouse primary somatosensory cortex. Neuron 61, 301–316 (2009).

31. D. H. O’Connor, S. P. Peron, D. Huber, K. Svoboda, Neural activity in barrel cortex underlying vibrissa-based object localization in mice. Neuron 67, 1048–1061 (2010).

32. F. Schnutgen et al., A directional strategy for monitoring Cre-mediated recombination at the cellular level in the mouse. Nat Biotechnol 21, 562–565 (2003).

33. D. Atasoy, Y. Aponte, H. H. Su, S. M. Sternson, A FLEX switch targets Channelrhodopsin-2 to multiple cell types for imaging and long-range circuit mapping. J Neurosci 28, 7025–7030 (2008).

34. D. van Versendaal et al., Elimination of inhibitory synapses is a major component of adult ocular dominance plasticity. Neuron 74, 374–383 (2012).

35. M. Fossati et al., SRGAP2 and Its Human-Specific Paralog Co-Regulate the Development of Excitatory and Inhibitory Synapses. Neuron 91, 356–369 (2016).

36. H. Chen et al., Fast assembling of neuron fragments in serial 3D sections. Brain Inform, (2017).

37. C. Charrier et al., Inhibition of SRGAP2 function by its human-specific paralogs induces neoteny during spine maturation. Cell 149, 923–935 (2012).

38. N. Spruston, Pyramidal neurons: dendritic structure and synaptic integration. Nat Rev Neurosci 9, 206–221 (2008).

39. P. Vetter, A. Roth, M. Hausser, Propagation of action potentials in dendrites depends on dendritic morphology. J Neurophysiol 85, 926–937 (2001).

40. M. Megias, Z. Emri, T. F. Freund, A. I. Gulyas, Total number and distribution of inhibitory and excitatory synapses on hippocampal CA1 pyramidal cells. Neuroscience 102, 527–540 (2001).

41. E. B. Bloss et al., Structured Dendritic Inhibition Supports Branch-Selective Integration in CA1 Pyramidal Cells. Neuron 89, 1016–1030 (2016).

42. L. Petreanu, T. Mao, S. M. Sternson, K. Svoboda, The subcellular organization of neocortical excitatory connections. Nature 457, 1142–1145 (2009).

43. T. Schikorski, C. F. Stevens, Morphological correlates of functionally defined synaptic vesicle populations. Nat Neurosci 4, 391–395 (2001).

44. F. Pennacchietti et al., Nanoscale Molecular Reorganization of the Inhibitory Postsynaptic Density Is a Determinant of GABAergic Synaptic Potentiation. J Neurosci 37, 1747–1756 (2017).

45. M. Matsuzaki, N. Honkura, G. C. Ellis-Davies, H. Kasai, Structural basis of long-term potentiation in single dendritic spines. Nature 429, 761–766 (2004).

46. E. M. Petrini et al., Synaptic recruitment of gephyrin regulates surface GABAA receptor dynamics for the expression of inhibitory LTP. Nat Commun 5, 3921 (2014).

47. A. Losonczy, J. K. Makara, J. C. Magee, Compartmentalized dendritic plasticity and input feature storage in neurons. Nature 452, 436–441 (2008).

48. B. Haider, D. A. McCormick, Rapid neocortical dynamics: cellular and network mechanisms. Neuron 62, 171–189 (2009).

49. J. S. Isaacson, M. Scanziani, How inhibition shapes cortical activity. Neuron 72, 231–243 (2011).

50. J. N. Bourne, K. M. Harris, Coordination of size and number of excitatory and inhibitory synapses results in a balanced structural plasticity along mature hippocampal CA1 dendrites during LTP. Hippocampus 21, 354–373 (2011).

51. H. Peng, Z. Ruan, F. Long, J. H. Simpson, E. W. Myers, V3D enables real-time 3D visualization and quantitative analysis of large-scale biological image data sets. Nat Biotechnol 28, 348–353 (2010).

52. H. Peng, A. Bria, Z. Zhou, G. Iannello, F. Long, Extensible visualization and analysis for multidimensional images using Vaa3D. Nat Protoc 9, 193–208 (2014).

53. A. Govindarajan, I. Israely, S. Y. Huang, S. Tonegawa, The dendritic branch is the preferred integrative unit for protein synthesis-dependent LTP. Neuron 69, 132–146 (2011).

54. S. L. Smith, I. T. Smith, T. Branco, M. Hausser, Dendritic spikes enhance stimulus selectivity in cortical neurons in vivo. Nature 503, 115–120 (2013).

55. N. Takahashi, T. G. Oertner, P. Hegemann, M. E. Larkum, Active cortical dendrites modulate perception. Science 354, 1587–1590 (2016).

56. N. L. Xu et al., Nonlinear dendritic integration of sensory and motor input during an active sensing task. Nature 492, 247–251 (2012).

57. P. Somogyi, G. Tamas, R. Lujan, E. H. Buhl, Salient features of synaptic organisation in the cerebral cortex. Brain Res Brain Res Rev 26, 113–135 (1998).

58. Y. Wang et al., Anatomical, physiological and molecular properties of Martinotti cells in the somatosensory cortex of the juvenile rat. J Physiol 561, 65–90 (2004).

59. R. C. Froemke, Plasticity of cortical excitatory-inhibitory balance. Annu Rev Neurosci 38, 195–219 (2015).

60. J. Waters, M. Larkum, B. Sakmann, F. Helmchen, Supralinear Ca2+ influx into dendritic tufts of layer 2/3 neocortical pyramidal neurons in vitro and in vivo. J Neurosci 23, 8558–8567 (2003).

61. R. Hand, F. Polleux, Neurogenin2 regulates the initial axon guidance of cortical pyramidal neurons projecting medially to the corpus callosum. Neural Dev 6, 30 (2011).

62. H. M. Carnevale NT, The NEURON Book. (Cambridge University Press, Cambridge, UK, 2006).

63. L. Sarid, R. Bruno, B. Sakmann, I. Segev, D. Feldmeyer, Modeling a layer 4-to-layer 2/3 module of a single column in rat neocortex: interweaving in vitro and in vivo experimental observations. Proc Natl Acad Sci U S A 104, 16353–16358 (2007).

64. H. Markram et al., Reconstruction and Simulation of Neocortical Microcircuitry. Cell 163, 456– 492 (2015).

65. A. Majewska, A. Tashiro, R. Yuste, Regulation of spine calcium dynamics by rapid spine motility. J Neurosci 20, 8262–8268 (2000).

66. Y. Kasugai et al., Quantitative localisation of synaptic and extrasynaptic GABAA receptor subunits on hippocampal pyramidal cells by freeze-fracture replica immunolabelling. Eur J Neurosci 32, 1868–1888 (2010).

67. C. Q. Chiu et al., Compartmentalization of GABAergic inhibition by dendritic spines. Science 340, 759–762 (2013).

68. G. A. Ascoli, Mobilizing the base of neuroscience data: the case of neuronal morphologies. Nat Rev Neurosci 7, 318–324 (2006).

69. L. Acciai, P. Soda, G. Iannello, Automated Neuron Tracing Methods: An Updated Account. Neuroinformatics 14, 353–367 (2016).

70. M. Halavi, K. A. Hamilton, R. Parekh, G. A. Ascoli, Digital reconstructions of neuronal morphology: three decades of research trends. Front Neurosci 6, 49 (2012).

71. E. Meijering, Neuron tracing in perspective. Cytometry A 77, 693–704 (2010).

72. A. Rodriguez, D. B. Ehlenberger, D. L. Dickstein, P. R. Hof, S. L. Wearne, Automated threedimensional detection and shape classification of dendritic spines from fluorescence microscopy images. PLoS One 3, e1997 (2008).

73. H. Peng, F. Long, G. Myers, Automatic 3D neuron tracing using all-path pruning. Bioinformatics 27, i239–247 (2011).

74. R. Barnes, C. Lehman, D. Mulla, Priority-flood: An optimal depression-filling and watershed-labeling algorithm for digital elevation models. Comput Geosci-Uk 62, 117–127 (2014).

